# Dynamic balance of myoplasmic energetics and redox state in a fast-twitch oxidative glycolytic skeletal muscle fiber

**DOI:** 10.1101/2025.07.03.662956

**Authors:** Disch Jana, A.L. Jeneson Jeroen, A. Beard Daniel, Röhrle Oliver, Klotz Thomas

## Abstract

In order to investigate the mechanisms governing energy and redox balance in skeletal muscle, we developed a computational model describing the coupled biochemical reaction network of glycolysis and mitochondrial oxidative phosphorylation (OxPhos) in fast-twitch oxidative glycolytic (FOG) muscle fibers. The model was identified against dynamic in vivo recordings of Phosphocreatine (PCr), inorganic Phosphate (Pi), and pH in rodent hindlimb muscle and verified against independent data from in vivo experiments and muscle biopsies.

Step response testing revealed that mass action kinetics in combination with feedback control were sufficient to accomplish myoplasmic ATP homeostasis over a 100-fold range of ATP turnover rates. This vital emergent property of the metabolic model was associated with dynamic behaviour of intermediary metabolite concentrations similar to a second-order underdamped system that remains to be verified. The simulations additionally predicted that the lactate dehydrogenase (LDH) reaction makes substantial contributions to redox balance across the physiological range of ATP demands in this myofiber phenotype, while its role in slowing cellular acidification is minimal. Yet, LDH knock-out simulations revealed that oxidative recycling of myoplasmic NADH in and by itself sufficed to maintain redox balance over ATP turnover rates in the range of mitochondrial ATP synthesis. We conclude that aerobic lactate production in working muscles is a byproduct of the metabolic flexibility of FOG myofibers afforded by expression of high levels of LDH and OxPhos enzymes to support continual myoplasmic redox balance and ATP synthesis under conditions of high-intensity mechanical work.

In the future, the presented simulation framework may be used to further enhance the understanding of how experimental observations in muscle emerge from the integrative behaviour of the metabolic network for carbohydrate metabolism in FOG myofibers.

## 1 Introduction

Skeletal muscles are typically composed of a mix of fatigue-resistant and fatigue-prone myofibers, affording performance of a wide range of mechanical tasks. Each myofiber phenotype is made up of a specific set of protein isoforms in concentrations tailored to the fiber’s particular function, ranging from slow-twitch (ST) oxidative ‘red’ fibers to fast-twitch (FT) glycogenolytic ‘white’ fibers (e.g. see [1, 2]). Fast-twitch oxidative glycolytic (FOG) myofibers represent a hybrid phenotype that ranks both functionally and metabolically in between these two fiber types.

The metabolic phenotypes of red and white myofibers represent opposite extremes, with red fibers reliant on the oxidation of fats and carbohydrates to fuel oxidative phosphorylation and white fibers principally relying on the fermentation of glycogen [1, 2], respectively, to maintain ATP balance during contractile work. The capillary contact surface areas of each of these two extreme myofiber phenotypes support these metabolic traits [3]. The intermediate FOG myofiber phenotype, on the other hand, has both a high concentration of glycolytic enzymes, including LDH, as well as a relatively high mitochondrial density and capillary contact surface [3]. As a result, FOG fibers can operate under conditions of both net pyruvate oxidation as well as fermentation. This ability endows FOG myofibers with metabolic flexibility to synthesize ATP under conditions of varying oxygen supply, such as may intermittently occur during any muscular work causing a collapse of capillaries due to increased intramuscular pressure [4].

This hybrid oxidative-anaerobic ATP synthetic metabolic network in FOG myofibers may, however, give rise to so-called ‘aerobic lactate production’ whereby a fraction of pyruvate is converted to myoplasmic lactate despite ample residual oxidative capacity [5, 6]. From a cellular metabolic efficiency viewpoint, any aerobic lactate production is wasteful since lactate can be readily lost to the extracellular milieu via any of the abundant organic anion exchangers in the myoplasmic membrane (e.g., MCT) [7], thus reducing the ATP yield per glucose by an order of magnitude. However, the net production of lactate from one cell type does not necessarily represent a metabolic inefficiency at the level of an organism, as lactate is used to supply reducing equivalents and pyruvate for oxidative ATP synthesis in the heart and hepatic gluconeogenesis during exercise. Over the 100-fold operational range of ATP turnover rates that skeletal FT myofibers must absorb and balance [8], the mechanisms governing net pyruvate mass flow into the mitochondrial reticulum versus myoplasmic LDH are important not only to redox and energy balance in the myofiber but also in the whole body.

In resting muscle, the cytoplasmic NAD^+^/NADH ratio has been estimated to be on the order of 400-900 [6]. Any production of myoplasmic NADH by GAPDH in the proximal part of glycolysis upon metabolic activation must be balanced by reoxidation of NAD^+^ to NADH by myoplasmic-mitochondrial NADH shuttles and/or LDH [6, 9]. Quantitative experimental methods to measure the dynamics of myoplasmic NAD(H) redox levels in situ are currently, however, missing. NADH fluorescence recordings from muscle, for example, are dominated by signal from mitochondrial matrix NADH [10].

To further understand the dynamic balance of myoplasmic energetics, redox state, and pH in an active fast-twitch muscle fiber, we used computer simulations to investigate ATP and pyruvate metabolism in FOG myofibers. First, we constructed a computational model of the coupled biochemical reaction network of glycolysis, LDH, and mitochondrial oxidative ADP phosphorylation (OxPhos). The model was next identified for the FOG myofiber phenotype against dynamic in vivo recordings of ATP metabolic levels and pH in rat FT muscle and validated against independent data from dynamic in vivo recordings in FT muscles as well as muscle biopsy analyses. Simulations then predicted that, on the one hand, mitochondrial pyruvate oxidation is the dominant source of ATP towards myoplasmic ATP balance over a 100-fold range of ATP turnover rates but myoplasmic NADH conversion by LDH, on the other hand, structurally contributes to myoplasmic redox balance and surpasses oxidation of NADH by mitochondrial shuttles about midway through this range.

## 2 Materials and methods

### 2.1 Model

The model describes energy metabolism in contracting fast-twitch oxidative glycolytic (FOG) fibers. The ATP turnover (Section 2.1.2) is balanced by ATP re-synthesis through (i.) the buffer reactions catalysed by creatine kinase (CK) and adenylate kinase (AK) (Section 2.1.2), (ii.) glyco(geno)lysis (Section 2.1.2), and (iii.) oxidative phosphorylation (OxPhos) (Section 2.1.2). The glycolytic and oxidative ADP phosphorylation are coupled to investigate proton and redox balance in the cytosol. This yields a network of biochemical reactions described through thermodynamically constrained kinetic equations. A schematic overview of the proposed model is shown in Fig. 1. The metabolic network is located in a myoplasmic compartment. Separate mitochondrial domains within the myoplasm were not included. Two additional compartments represent the interstitial fluid and capillaries, respectively (see Supplementary Fig. 1).

**Fig 1.**
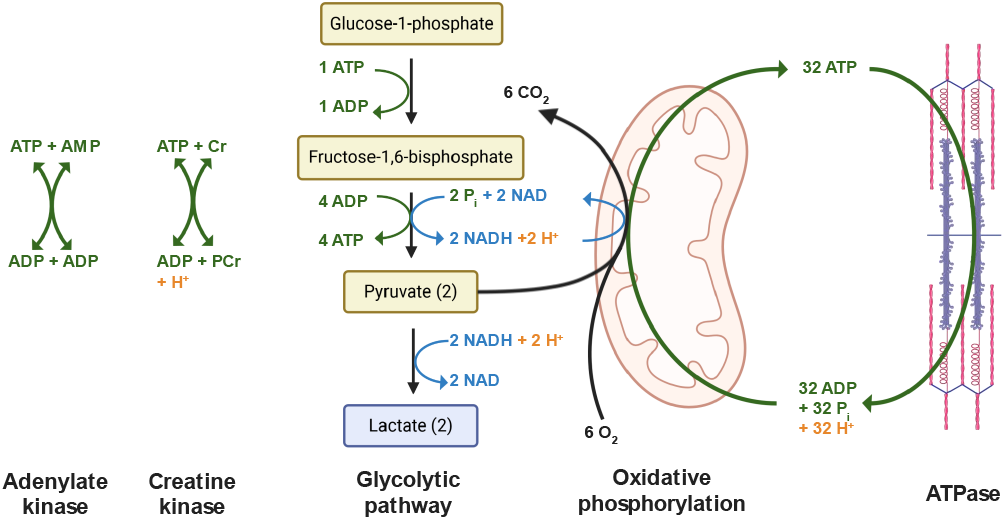
Schematic drawing of the coupled biochemical network involved in the dynamic balance of ATP (green), redox state (blue), and pH dynamics (red) in skeletal muscle fibers. This figure was created in BioRender.com.

#### 2.1.1 Basic concept

In this section, we provide a general description of the utilized modeling framework. For further details, see [11, 12].

##### Proton balance

Most metabolites are (negatively charged) ions and, hence, exist in various species (bound to protons or metal ions), with rapid kinetic transitions that can be assumed to be in thermodynamic equilibrium. The total concentration of an arbitrary metabolite L is thus given by

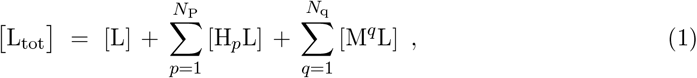

with [L] representing the concentration of the unbound, most deprotonated species in a pH range of 5.5–8.5. Further, *p* denotes the stoichiometry number of bound protons with *N*_P_ denoting the number of protonised species and *N*_q_ is the number of metal-bound species. As a simplification, we assume that only one metal ion of each type can bind to L (with *N*_q_ = 2 whereby M^1^ = Mg^2+^ and M^2^ = K^+^) and proton binding to metal-bound species is neglected [12]. The pH-dependent concentration of each species can be computed using the dissociation constants:

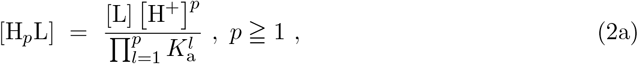

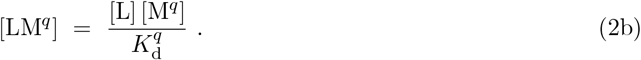

Therein, 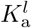 is the acid dissociation constant of the reaction [H_*l*_L] ⇄ [H_*l*−1_L] + H^+^ and 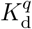 the dissociation constant for the binding reaction to the given metal ion type M^*q*^. Thereby, the dissociation constants 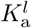 are empirically corrected to the simulated temperature and ionic strength.

Using Eqn. (2), one can compute the average number of protons bound to metabolite L:

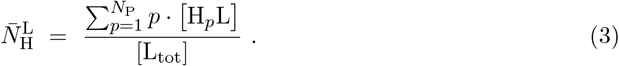

Thus, for an arbitrary biochemical reaction *k* one obtains the following proton consumption stoichiometry

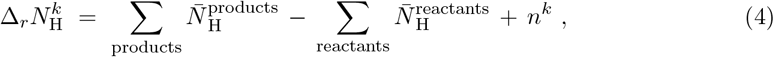

with *n*^*k*^ denoting the number of protons consumed by the associated reference reaction. In the proposed model, all reference reactions are defined in terms of the most deprotonated species in the pH range of 5.5–8.5 (see Supplementary Table 1) and are balanced with respect to mass and charge [12]. The proton consumption flux of a biochemical reaction is the product of the flux through the (reversible) biochemical reaction *J*^*k*^ and the proton consumption stoichiometry 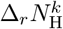:

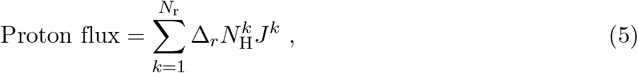

where *N*_r_ denotes the total number of reactions in the network. Note that changes in proton concentration and thus pH (pH = − log_10_([H^+^])) have, in turn, an effect on the reaction fluxes *J*^*k*^ since both the maximal velocity of a biochemical reaction, 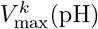, and the apparent equilibrium constant, 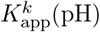, are a function of pH as described in the following sections.

##### Kinetics

The enzyme-catalyzed reaction fluxes *J*^*k*^ are (unless specified differently) defined as reversible reactions based on Michaelis-Menten kinetics, i.e.,

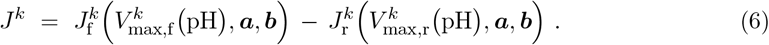

Therein, 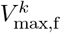 and 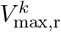 denote the maximal reaction velocity in forward and reverse direction, respectively. For enzyme reactions with known pH optimum, *V*_max,f_ is described using an empirical pH correction [12]. Further, ***a*** is a vector of substrate and product concentrations and ***b*** contains the kinetic (Michaelis) constants for the substrates (S) and products (P).

##### Thermodynamics

The reversible reaction kinetics in the model are thermodynamically constrained through the Haldane relation, i.e., relating the apparent equilibrium constant, 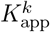, and the kinetic parameters:

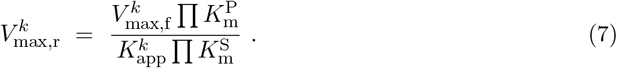

Therein 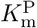 and 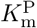 denote the Michaelis constants of substrates and products respectively. The apparent equilibrium constant 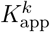 of reaction *k* is defined in terms of the equilibrium constant of the associated reference reaction, 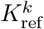, using the so-called binding polynomials (see [11]):

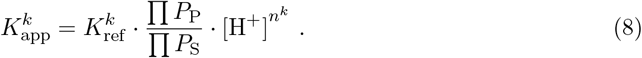

The binding polynomials for the substrates (*P*_S_) and products (*P*_P_) of reaction *k* are given by

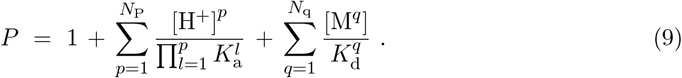

Further, the equilibrium constant of each reference reaction in the model is calculated using the standard free energy of the reference reaction 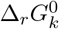, i.e., the difference between the free energies of the formation of the products and reactants:

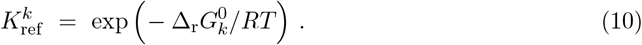

Therein, *R* =8.315 J K^−1^ mol^−1^ is the universal gas constant and *T* is the absolute temperature in Kelvin. The transformed Gibbs energy of each biochemical reaction can be calculated as

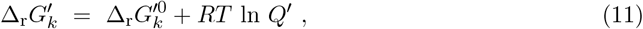

where the *Q*^*′*^ is the apparent reaction quotient, i.e. the reaction quotient of the biochemical reaction defined in terms of the sums of species. Further, the standard transformed Gibbs energy, 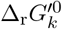, can be derived from the apparent equilibrium constant (see Eqn. (8)):

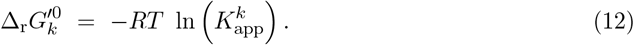

#### 2.1.2 Muscle fiber model

This section provides a brief description of the metabolic model describing an FOG muscle fiber. A detailed description of each reaction is given in Appendix B.1.3.

##### ATPase activity

We use a lumped total ATP consumption flux (e.g., cross-bridge cycling and SERCA pumps) that is modeled as the sum of a basal and an activity-dependent rate. The basal rate is fixed at 0.48 mM*/*min as observed in resting skeletal muscle [13]. The activation rate is modeled as a step function with variable amplitude and time intervals.

##### ATP buffers

Creatine kinase (CK) catalyses the reaction PCr + ADP ⇄ Cr + ATP, that serves as an instantaneous ATP buffer in response to changes in the ATPase activity. For the CK reaction, the free energy of formation of the reference species HCr^0^ and HPCr^2-^ (cf. Supplementary Table 1) were not available. Thus, instead of calculating the equilibrium constant as described in Eqn. (10), the apparent equilibrium constant is given by

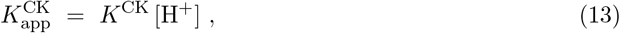

where *K*^CK^ = 1.66 *×* 10^9^ M^−1^ is a reported literature value [14]. Note that the reported *K*^CK^ was determined at 38 ^°^C, pH = 7, an ionic strength of 0.25 M and an Mg^2+^ concentration of 1 mM, whereby the effect of potassium binding was not considered. These conditions slightly deviate from the proposed model with a temperature of 37 ^°^C and an ionic strength of I = 0.1 M. Further, the model captures ATP buffering through Adenylate kinase (AK).

##### Glycolysis

During rest-to-work transitions, glycolytic ATP synthesis rapidly increases, and glycolysis is particularly relevant during high-intensity exercise. In this work, we describe the biochemical reaction network responsible for the turnover of glucose-1-phosphate (G1P) to pyruvate (PYR) and lactate (LAC) through various glycolytic intermediates (see Table 1). Thereby, Phosphofructokinase (PFK) is known to be a key regulatory enzyme that facilitates the rapid control of the glycolytic flux. We used a pseudorandom-order kinetic PFK model with statistical inhibition developed by Waser et al. [15] and later adopted by Connett [16, 17]. The PFK model considers activation due to an alkaline shift in cytoplasmic pH as well as ATP mediated inhibition and competitive deinhibition by the metabolites ADP and AMP. Thereby, the binding constants for ADP and AMP (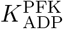 and 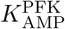, see Table 2) were adjusted heuristically to achieve a glycolytic flux near zero in the resting state while simultaneously facilitating a rapid glycolytic flux regulation when the muscle is activated. This behaviour closely mimics experimental observations in fast-twitch muscles [18–21].

**Table 1.** Maximal enzyme activities.

| Enzyme | Abbreviation | $V_{\max}$ (M/min) |
| --- | --- | --- |
| Phosphoglucomutase | PGLM | 2.7<sup>b</sup> |
| Phosphoglucoisomerase | PGI | 1.2<sup>a,*</sup> |
| Phosphoglucokinase | PFK | 0.13<sup>b</sup> |
| Aldolase | ALD | 0.2<sup>a</sup> |
| Triosephosphate isomerase | TPI | 24.0<sup>a</sup> |
| Glycol-3-phosphate dehydrogenase | G3PDH | 1.3<sup>b,*</sup> |
| Glyceraldehyd-3-phosphate dehydrogenase | GAPDH | 3.3<sup>a</sup> |
| Phosphoglycerate kinase | PGK | 2.0<sup>a,*</sup> |
| Phosphoglycerate mutase | PGM | 1.6<sup>a</sup> |
| Enolase | ENO | 0.3<sup>a</sup> |
| Pyruvate kinase | PYK | 0.2<sup>b</sup> |
| Lactate dehydrogenase | LDH | 4.0<sup>a</sup> |
| Adenylate kinase | AK | 1.2<sup>a</sup> |
| Creatine kinase | CK | 1.0<sup>a,*</sup> |

<sup>a</sup>: Reported in [34]; <sup>b</sup>: heuristically adjusted; <sup>\*</sup>: Given as reverse rate.

**Table 2.** optimized parameters.

| Parameter | Description | Value | CoV (%) |
| --- | --- | --- | --- |
| $V_{\max}^{\text{OxPhos}}$ | Maximal reaction velocity of OxPhos | 1.25 mM/min | 1.1 |
| $n_{\text{H}}$ | Hill coefficient for ADP of OxPhos | 3.0 (-) | 4.0 |
| $K_{\text{ADP}}^{\text{OxPhos}}$ | Half saturation concentration of ADP for OxPhos | 30.121 $\mu\text{M}$ | 10.4 |
| $[\text{X}]_{\text{T}}$ | Total concentration of intrinsic buffer<br>(non-phosphate, non-carbonate) | 79.7 mM | 5.5 |
| TPP | Total phosphate pool | 60.2 mM | 1.2 |
| $V_{\text{act}}^{\text{ATPase}}(2 \text{ Hz})$ | ATPase rate for 2 Hz stimulation | 22.5 mM/min | 4.4 |
| $V_{\text{act}}^{\text{ATPase}}(4 \text{ Hz})$ | ATPase rate for 4 Hz stimulation | 40.5 mM/min | 2.3 |
| $V_{\text{act}}^{\text{ATPase}}(10 \text{ Hz})$ | ATPase rate for 10 Hz stimulation | 50.3 mM/min | 2.0 |

Note that we neglected glycogen phosphorylase (both isoforms A and B), i.e., converting glycogen into G1P, as there exists no kinetic model adequately describing the dynamics of this reaction. Instead, the G1P concentration was clamped ([G1P] = 0.07 mM), acting as a substrate pool for phosphoglucomutase (PGLM). Theoretically, this would yield an increasing total phosphate pool (TPP). Thus, we modified the time derivative of inorganic phosphate to include the PGLM flux in a way that Pi is removed from the myoplasm at the rate of G1P turnover via PGLM (see Appendix B.2).

##### Mitochondrial ATP synthesis

Oxidative phosphorylation in the mitochondria is the most efficient and sustainable ATP synthesis pathway. Although mitochondria can use different fuels (e.g., fatty acids), given that we are modeling FOG fibers, we only considered oxidative carbohydrate metabolism [22]. A top-down approach was used, where all biochemical reactions associated with mitochondria, i.e., pyruvate dehydrogenase, TCA cycle (tricarboxylic acid cycle), electron transport chain, and ATP synthase, are lumped together in one reaction (see Appendix B.1.5). This yields the following reference reaction:

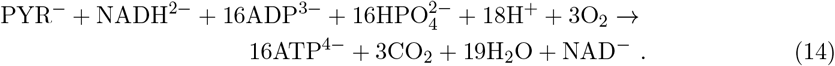

Equation (14) considers key intermediates and substrates that couple the glycolytic and aerobic pathways. Note that with limited availability of myoplasmic NADH, the capacity for ATP production by the OxPhos model is reduced, which is described in Appendix B.1.5.

The kinetics of OxPhos model is based on a sigmoid Hill function with feedback regulation of the substrates ADP [23], inorganic phosphate (Pi), and pyruvate, i.e.,

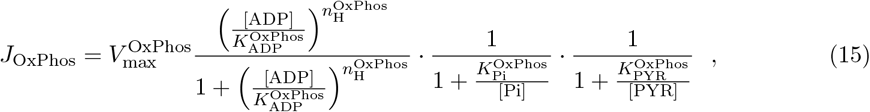

where the constants 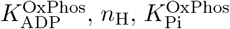 and 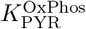 determine the sensitivity to changes of the substrate concentrations. In detail, 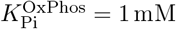 is the half-saturation concentration of Pi, whereby reported *K* _m_ values from in vitro observations in purified mammalian heart and liver mitochondria range from 0.25 mM to 1 mM [24, 25]. Further, 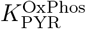 was set to 0.15 mM, a value reported to be the *K* _m_ for pyruvate translocation into rat liver mitochondria [26]. This value is within the range of other reported *K* _m_ values for pyruvate [27–30]. Lastly, the parameters 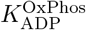, as well as the maximal reaction velocity, 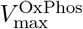, and the Hill coefficient 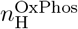 were optimized using dynamic phosphorus magnetic resonance spectroscopy data as described in Section 2.2.2.

##### Generic proton buffer

In addition to the dynamic proton buffer capacity of the metabolites, the model also considers buffering via e.g. proteins by assuming a constant intrinsic buffer size [X]_T_ with a dissociation constant of *K*_*a*_ = 1 *×* 10^−7^ M for a working range around pH = 7. The buffer concentration [X]_T_ is the sum of the acetic acid [HX] and its conjugate base [X^-^]. The dissociation constant *K* _a_ is defined as *K* _a_ = [H^+^][X^-^]/[HX]. Moreover, the buffer capacity *β* is defined as the concentration of acid (*C* _a_) or base (*C* _b_) that needs to be added to a buffer solution to change its pH by one unit (in slykes i.e. mmol/L/pH unit). In a physiological pH range, the buffer concentration of a weak monoprotonic acid can be calculated as follows [31]:

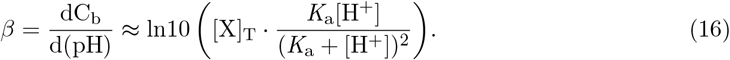

##### Metabolite washout

Energy metabolism in skeletal muscles requires product/substrate exchange with the environment. To simulate an open system, the model is split up into three compartments representing the muscle fiber, the extracellular fluid, and the capillaries. The compartments are coupled through metabolite-specific transport systems. To couple the muscle fiber and the extracellular environment, we consider passive and facilitated transmembrane transport of lactate, CO_2_, and bicarbonate. Lactate-proton symport is achieved by the monocarboxylate transporter MCT, where we only considered isoform MCT4 (i.e., the predominant isoform in cells with a high glycolytic rate [32]). Note that transport via MCT is bi-directional, allowing the utilization of extracellular lactate generated by other cells as a fuel for the aerobic pathway. This is known as the intercellular lactate shuttle [33]. In this work, extracellular lactate levels were clamped to 1 mM. The transport of CO_2_ over the muscle fiber membrane occurs by passive diffusion. Further, CO_2_ and bicarbonate can be transported from the extracellular space into the capillary compartment. The kinetic equations and parameters of all transport reactions are provided in Appendix B.1.7.

#### 2.1.3 Parameters and initial conditions

The model consists of 85 adjustable parameters (excluding the 13 apparent equilibrium constants which are dynamically computed as described in Section 2.1.1) and 41 state variables. Most parameters could be fixed based on existing data from the literature. Only 5 parameters were identified by solving an optimization problem (Section 2.2.2).

##### Enzyme activities

Maximum enzyme activities of the glycolytic reaction network, as well as CK and AK, measured at 37 ^°^C in rabbit skeletal muscle are reported in [34]. Four maximum enzyme activity values (PGLM, PFK, G3PDH, and PYK) needed to be adjusted heuristically to handle a large range of ATP turnover rates without excessive accumulation of glycolytic intermediates under normal physiological conditions. Table 1 summarizes the maximal activities for each enzyme-catalyzed reaction.

##### Baseline concentrations

In order to obtain steady-state baseline concentrations that are consistent with experimental data from fast twitch muscles, we set the initial concentrations of the metabolites in the model to appropriate values, primarily focusing on intracellular ATP, ADP, PCr, Pi and pH as well as intracellular lactate, since the baseline concentrations of other metabolites such as the glycolytic intermediates cannot be measured in an intact physiological system. The system is constrained by mass conservation where the total creatine pool sums to 45.2 mM and the sum of total adenine nucleotides is 10.32 mM. The size of the total phosphate pool (TPP) is defined in terms of transferable phosphates and was fitted to experimental data as described in Section 2.2.2). Using the guessed initial concentrations, we conducted simulations at the basal ATP demand until the model reached a steady state as part of the model identification (see Section 2.2.2). The resulting steady-state baseline concentrations of the model (given the optimized parametrization) can be found in Supplementary Table 4.

##### Compartments and volume fractions

The volume fractions 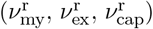 and the water content 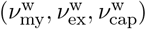 of the model compartments myoplasm, extracellular space, and capillaries are needed as a scaling factor in the differential equations of each metabolite (see Appendix B.2). Their values were derived using data from frog muscle [35]. In short, the total water content of 1 g wet-weight frog muscle (sartorius) is 78.5 %, whereby 12.5 % can be attributed to the extracellular space and 66 % to the muscle fiber. Further, we used the simplified assumption that the extracellular and capillary compartments consist of 100 % water, i.e., 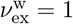 and 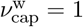. For the myoplasm compartment, this leads to a volume fraction of 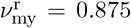 and water content 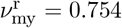. The volume fractions of the interstitial and cytoplasmic compartments are assumed to be 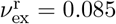 and 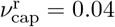, respectively.

### 2.2 Simulations

#### 2.2.1 Computer model

The model described in Section 2.1 yields a differential-algebraic system of equations (DAE). A numerical model was implemented using MATLAB 9.12 (Mathworks Inc., MA, USA). To obtain the right-hand side of the system of differential equations describing the biochemical reaction network, we used the MATLAB-based Biochemical Simulation Environment (BISEN, Medical College of Wisconsin, WI, USA) [36]. The differential equations were solved numerically using MATLAB’s built-in ‘ode15s’ solver optimized to handle stiff initial value problems (absolute and relative tolerance: 10^-9^, maximal step size: 3 s). Thereby, it is required that the metabolite concentrations are non-negative.

#### 2.2.2 Model identification

A set of 5 parameters (i.e., the maximum OxPhos rate, the Hill coefficient of the OxPhos model, the sensitivity of OxPhos to the ADP concentration, the generic buffer size and the total phosphate pool) and the ATPase activity corresponding to a specific experiment were identified by fitting the model to experimental data published by Kushmerick et al. [37]. In the following, ***q*** denotes the vector containing the model parameters and initial conditions to be estimated by solving an optimization problem.

##### Identification data

The data used for the model identification was collected from the gastrocnemius-plantaris muscle of anesthetized rats via ^31^P magnetic resonance spectroscopy (^31^P-MRS). Kushmerick et al. [37] acquired dynamic ^31^P-MRS data before, during, and after muscle activation. The muscle was activated by electrically stimulating the sciatic nerve at frequencies of 2 Hz, 4 Hz, and 10 Hz. For each stimulation frequency, the time course of [PCr], [Pi], and intracellular pH was derived from the relative peak areas of the collected spectra.

##### Optimization problem

To quantify the difference between model predictions and experimental data, a fitness function *L*(***q***) was defined as the weighted mean absolute error (MAE) between the (regularized) experimental data ***x***(*t*_*i*_) (with *t*_*i*_ denoting the temporal sampling points and *i* = 1, …, *n*_i_) and corresponding simulated observations 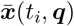. The regularization was introduced as ^31^P-MRS quantifies relative changes in metabolite concentration. Thus, we compared the relative change of the simulated and measured concentrations. Thereby, approximately 1 mM of the measured Pi is considered to belong to the extracellular environment [37]. This was taken into account by adding 1 mM Pi to the simulated time course of intracellular Pi before calculating the error between the model simulation and experimental data. Further, as the simulated baseline pH depends on the selected parameters, we compared the absolute change of the pH values with ΔpH(***q***) denoting the difference of the baseline pH between the experimental data and the simulation. This yields

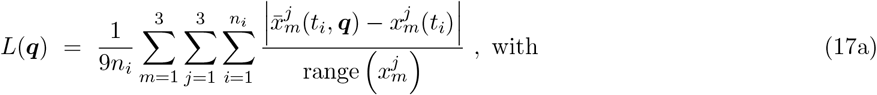

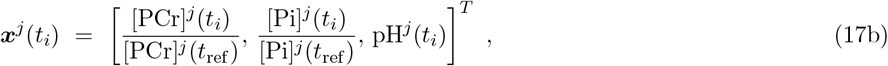

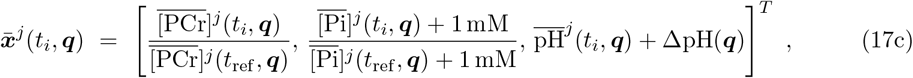

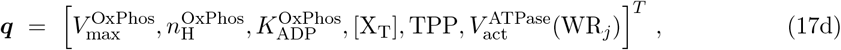

where the indices *m* refer to the measured variable (*m* = 1: PCr, *m* = 2: Pi, *m* = 3: pH) and *j* denotes the stimulation frequency (*j* = 1: 2 Hz, *j* = 2: 4 Hz, *j* = 3: 10 Hz). Further, the reciprocal of the range function (i.e., the maximum minus the minimum value) of the individual time series data were used as weights.

Equation (17a) was minimized using a genetic algorithm provided by MATLAB’s global optimization toolbox and the function ‘ga’ (default settings). Each population contains 200 parameter sets, with the output of the optimization problem being the parameter set with the minimum loss in the last generation. The genetic optimization iteratively updates the population of parameters considering (quasi-)random mutations. Hence, the obtained solution depends on the seed of the utilized random number generator as well as the initial population. To test the uniqueness of the estimated parameters, we ran the optimization with 20 different first generations. Finally, we computed the coefficient of variation for each parameter as well as the linear correlation between all sets of parameters.

In order to fit the model to the experimental data, one simulation was performed per stimulation frequency following a protocol of 1.8 minutes of muscle activation with an ATPase rate above basal level and a subsequent 3 min long recovery phase. An initialization phase at the basal ATPase rate was implemented to ensure that the model has reached a steady-state baseline with each parameter set ***q*** before the simulation protocol begins. The ATP turnover rates during muscle activation were estimated by the optimization problem (see Table 2).

##### Physiological constraints

We restricted the solution space of the estimated parameters ***q*** by specifying lower and upper bounds corresponding to reported values from the literature (if available). The Hill coefficient 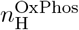 of the OxPhos model (Eqn. (15)) was found to be at least 2 with values reported not higher than 3 [23, 38, 39]. For isolated mitochondria extracted from mammalian heart muscle or liver, reported *K* _m_ values for ADP have been typically on the order of 30 µM [25, 40]. Thus, the permissible parameter range of 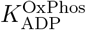 was chosen between 10 µM and 100 µM. The maximal oxidative ATP synthesis rate was computed based on the maximal increase in oxygen consumption during tetanic contraction of 3.62 µmol O_2_ min^-1^ g muscle^-1^ observed in perfused rat hindlimb [41]. Using ATP/O_2_ = 5.33 (Eqn. (14)) and a factor of 0.66 [35] to account for the fraction of intracellular water per unit wet weight of muscle, the *V* 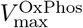 value was calculated to be approximately 30/16 mM/min. However, the upper bound for 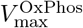 was lowered to 20/16 mM/min in order to prevent oscillations that occurred during simulations with low ATPase rates caused by the coupled feedback regulation of the glycolytic flux and OxPhos.

The protein buffer capacity of human muscle is estimated to be 15-45 slykes plus an additional contribution of around 3 slykes by dipeptide anserine [42]. This yields a lower and upper bound for the buffer concentration of [X]_T,lower_ = 0.031 M and [X]_T,upper_ = 0.084 M respectively given a basal pH value of around 7.05 (Eqn. (16)).

Note that the design variables have an effect on the steady-state baseline levels of the metabolites. Yet, the ratio of PCr and Pi in the model and the experimental data needs to be similar to ensure comparability. Thus, the total phosphate pool (TPP) was optimized by constraining the initial Pi concentration between 0 and 4 mM. This yields a permissible TPP range from

#### 2.2.3 Model validation

The model was validated against independent in vivo ^31^P-MRS data of intracellular pH and PCr collected by Foley et al. [43] during rest, muscle contraction, and recovery. Gastrocnemius muscles of anesthetized rats were stimulated at 0.75 Hz for 8 minutes. In contrast to the stimulation frequencies used for model identification, muscle stimulations at 0.75 Hz are below the mitochondrial ATP synthesis capacity (at approx. 0.8 Hz [38]). The ATPase rate, i.e., the model input, corresponding to a frequency of 0.75 Hz was calculated from the initial rate of PCr hydrolysis, assuming that neither glycolysis nor oxidative phosphorylation has a significant contribution to ATP supply right at the onset of muscle stimulation. The ATPase rate was reported to be (11.51 *±* 1.47) µmol g^−1^ min^−1^ calculated from a monoexponential fit to the measured PCr changes [43]. Using a factor of 0.66 [35] to account for the fraction of intracellular water per unit wet weight of muscle, the calculated ATPase rate is (17.44 *±* 2.23) mmol L^−1^ min^−1^, i.e., below the model’s maximal capacity for oxidative ATP synthesis of 20 mmol L^−1^ min^−1^.

We also estimated the ATPase rate during exercise by fitting the model output directly to the measured PCr dynamics [43] using a genetic algorithm and the same method as for the parameter identification described in the section above. Note that in this case, there is only one stimulation frequency (j = 1: 0.75 Hz) and one measured variable (m = 1: PCr) considered in *L*(***q***) and consequently ***x***^*j*^(*t*_*i*_) and ***x***^*j*^(*t*_*i*_, ***q***). Thereby, the ATPase rate was the only design variable in ***q***. The ATPase rate during the exercise was determined to be 11.24 mmol L^−1^ min^−1^.

#### 2.2.4 Multi parametric sensitivity analysis

A local multi-parameter sensitivity analysis (MPSA) [44–47] was performed to evaluate the sensitivity of the model to (simultaneous) variations of kinetic parameters and the total phosphate, creatine, adenosine, and redox pool sizes as well as the intrinsic proton buffer size. In short, the MPSA is based on a Monte Carlo method in which the model is run repeatedly using different parameter sets drawn from a uniform probability distribution for each tested parameter. All parameters were varied by *±*1 %. The sensitivity values *D*_*m,n*_ *∈* [0, 1] are computed using the Kolmogorov-Smirnov statistic and quantify the influence of variations of a parameter *n* on a state variable *m*, with sensitivity values close to one indicating high sensitivities. A detailed step-by-step description of the performed MPSA can be found in Appendix C.

#### 2.2.5 Model predictions

##### Experiment 1 – Intracellular redox potential

Currently, no in vivo modality can directly measure internal state variables, such as the concentration of glycolytic intermediates or the cytoplasmic redox potential. We use the proposed computational model to predict the internal dynamics of the given biochemical network. For this purpose, we simulated 10 min of muscle activation with variable contraction intensities, i.e., from 0.5 mM*/*min to 30 mM*/*min ATPase rates with a step size of 0.5 mM*/*min, followed by a recovery phase with an ATPase rate at the basal level.

##### Experiment 2 – LDH-KO

Lactate dehydrogenase (LDH) catalyzes the reversible reduction of pyruvate to lactate while simultaneously oxidizing NADH. To study the role of LDH in ATP homeostasis and redox balance, we build an LDH knockout model (i.e., 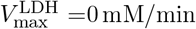). We simulated a 20 min long exercise protocol at different ATPase rates between 0.5 mM*/*min and 20 mM*/*min, i.e. the maximal capacity of oxidative ATP production, with a step size of 0.25 mM*/*min. Since the knock-out of LDH influences the basal state of the muscle fiber, the knock-out model’s resting state was determined by running the model without applying a stimulus prior to the simulation protocol. During the initialization phase, the pH value was clamped since the results from the MPSA indicate a strong influence of pH on the model baseline (see Section 3.1.3). For comparison, we applied the same exercise protocols to the control model with normal LDH activity (i.e.,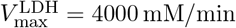).

##### Experiment 3 – LDH-KO under ischaemia

Further, we tested the effects of perturbations in myoplasmic redox balance on ATP metabolism caused by knocked out LDH under ischaemic conditions. Ischemia was simulated by setting 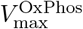 to 0 mM*/*min at the onset of muscle activation. In addition, the removal of lactate and carbon dioxide from the extracellular space was prevented by unclamping extracellular lactate and setting transport mechanisms from the extracellular to the capillary compartment to zero. We compared three different model configurations, i.e., the (healthy) control model, the LDH KO model, and an LDH KO model with clamped NAD^+^ and NADH concentrations, when prescribing an ATPase rate at of 20 mM*/*min with a duration of 1 min.

**Supplementary Table 2.**
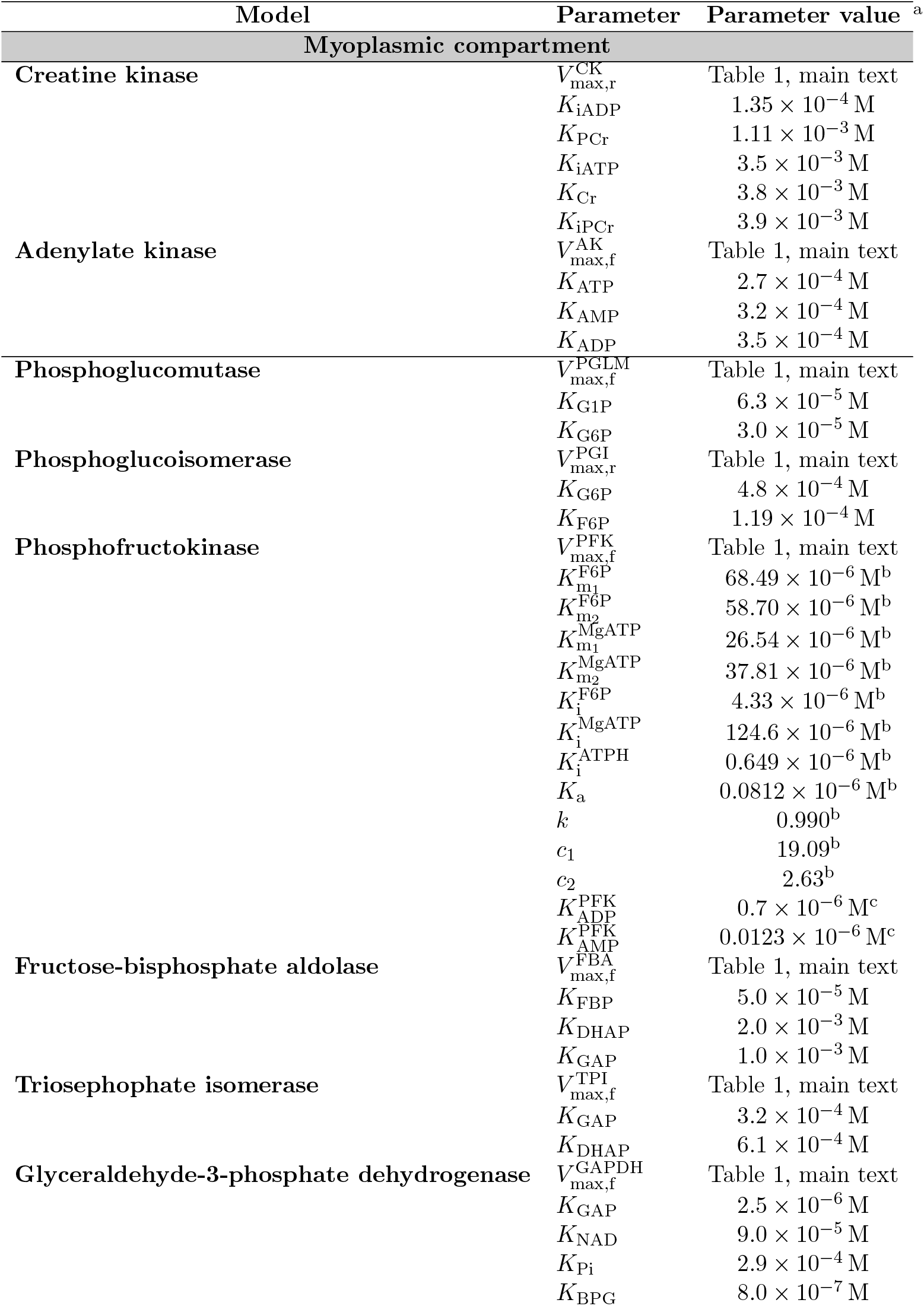
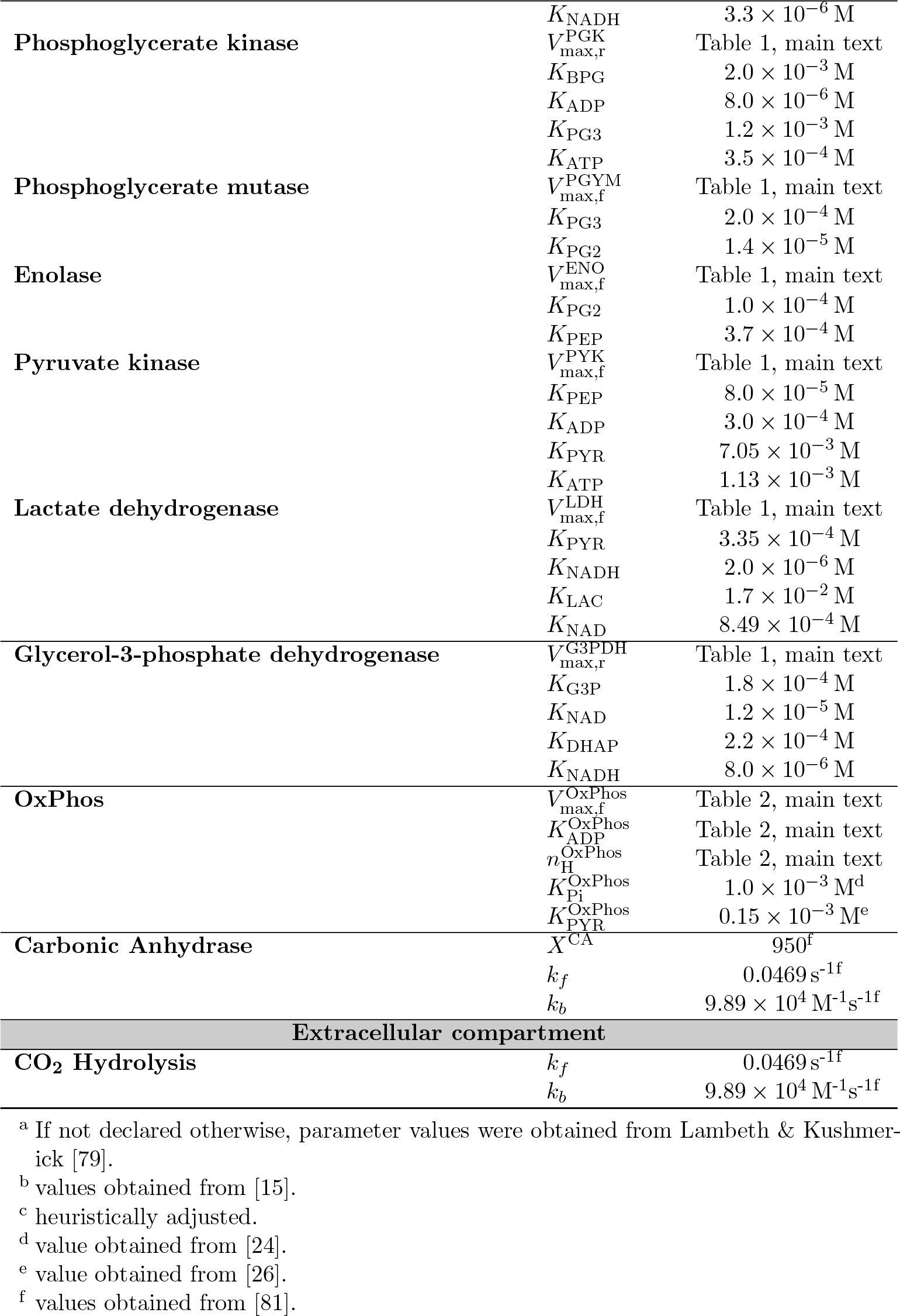
Kinetic Parameters of the biochemical reaction models.

## 3 Results

### 3.1 Model development

#### 3.1.1 Parameter Identification

A few model parameters that could not be determined through literature values were fitted to experimental data [37] using a genetic optimization algorithm (see Section 2.2.2). The optimized parameter set with the smallest loss function value out of the 20 optimization runs was used for the parametrization of the model. The parameter values are given in Table 2 together with the coefficient of variation (CoV) computed across the 20 optimization runs. Notably, the CoV is not bigger than 10.4 % for all estimated parameters, demonstrating the robustness of the parameter identification method.

Fig. 2 compares the pH, Pi, and PCr dynamics of the fitted model and experimental data. It can be observed that the model accurately captures the measured dynamics of PCr, Pi, and pH. The simulated PCr concentration decreases by 56 %, 83 % and 91 % during muscle activations of 2 Hz, 4 Hz and 10 Hz, respectively. These values slightly exceed the in vivo data for all three stimulation frequencies. We note that for the 4 Hz and 10 Hz stimulation frequency, the initial drop in PCr is well matched by the simulation. However, the experimental data of PCr shows a recovery of PCr during the second half of muscle activation (see Fig. 2, panels B and C) that is not captured by the model. Due to the coupling of ATP hydrolysis and resynthesis through the creatine kinase reaction, the Pi and PCr dynamics show a strong negative correlation. The simulations show a fivefold, sevenfold, and eightfold increase in Pi during muscle activations for the 2 Hz, 4 Hz, and 10 Hz simulations, respectively. For the 2 Hz simulation, the in silico predictions match the in vivo data quantitatively (MAE = 1.6 mM). However, the model underestimates the increase in Pi for the 4 Hz simulation (MAE = 3.2 mM) as well as the amplitude of the dynamic Pi response for the 10 Hz simulation (MAE = 4.3 mM). Notably, the measured time courses of PCr and Pi show a bump that is also visible in the simulation results. The model also captures the magnitude of change in pH with the initial alkalinization during ATP supply through the creatine kinase reaction, followed by acidification caused by the activation of the glycolytic pathway. Yet, the measured acidification exceeds the simulated pH drop for all three stimulation frequencies.

**Fig 2.**
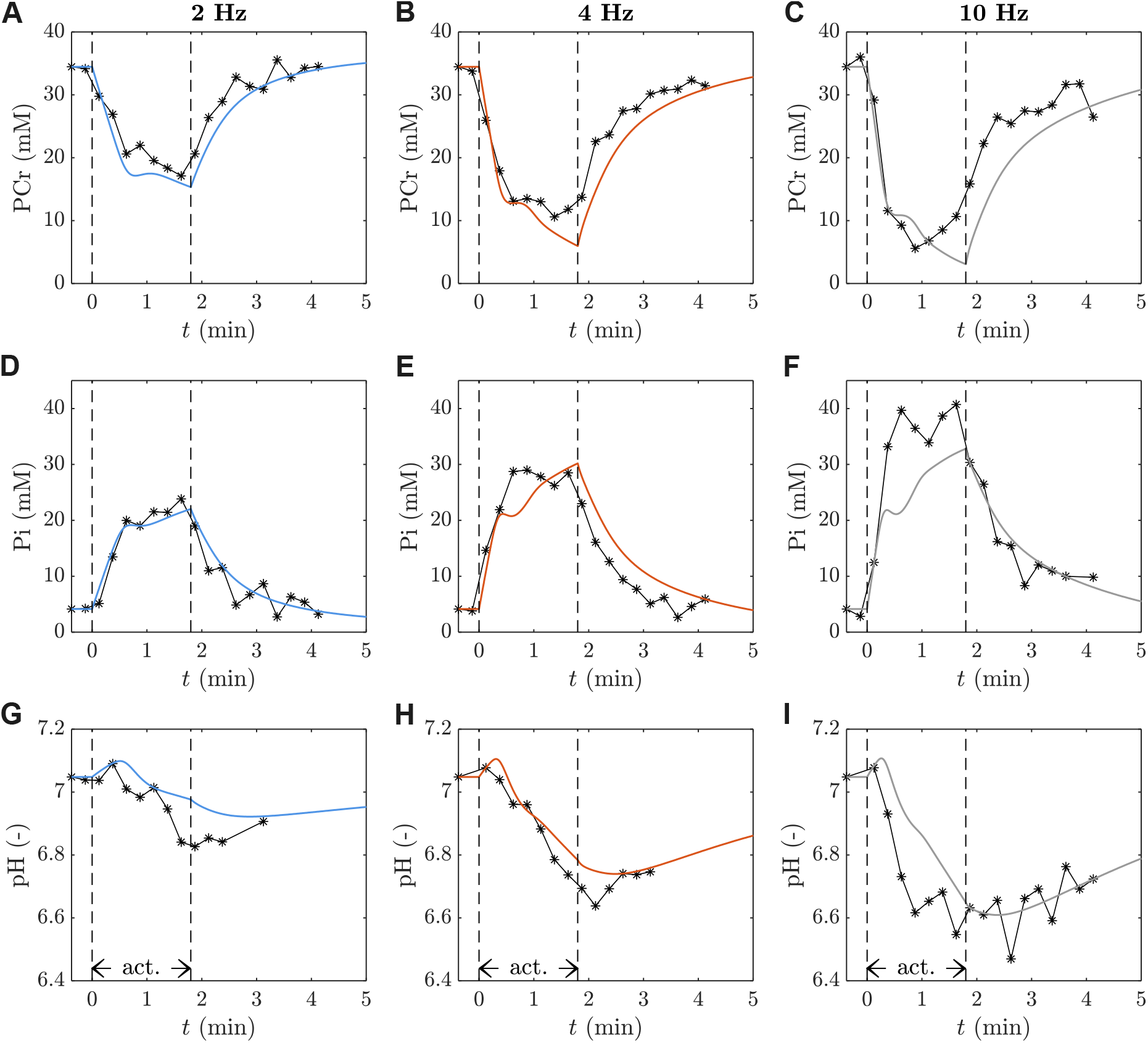
Comparison between the optimized model output (solid lines) and measured [37] (markers) [PCr] (A-C), [Pi] (D-F), and pH dynamics (G-I), depending on the stimulation frequency (Left: 2 Hz; Middle: 4 Hz; Right: 10 Hz). Note that the visualized data is normalized such that the simulated and measured data have the same baseline, see Section 2.2.2.

Lastly, we computed the correlation between the estimated parameters (see Supplementary Fig. 2). The ATPase rates at 4 Hz and 10 Hz show a strong positive correlation with the generic proton buffer capacity [X]_T_ (*r* ≧ 0.89; *p* ≪ 0.001). This is accompanied by a strong positive correlation between the ATPase rates at 4 Hz and 10 Hz (*r* ≧ 0.75; *p* ≪ 0.001). Further, a strong negative correlation is detected between the total phosphate pool and 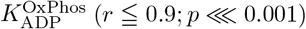. Finally, we note that the values of 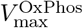 and 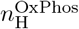 are equal to the nprescribed upper bound values.

#### 3.1.2 Model verification

To verify the metabolic muscle fiber model, in silico predictions were compared to independent in vivo ^31^P-MRS data of pH and PCr dynamics collected by Foley et al. [43] (see Fig. 3). The experiment uses an 8 min long electric stimulation protocol with a stimulation frequency of 0.75 Hz. The corresponding ATP hydrolysis rate was given as (17.44 *±* 2.23) mM*/*min [43] and derived from the time derivative of the measured PCr concentration, i.e., approximated by a mono-exponential fit, at the onset of stimulation. The results visualized in Fig. 3 show that the model qualitatively reproduces PCr and pH dynamics given an ATP hydrolysis rate outside the model’s identification range. However, the model overestimates both the drop in PCr concentration and pH. Treating the ATPase rate as an adjustable parameter optimized with a genetic algorithm yielded a lower ATPase rate of 11.28 mM*/*min (Fig. 3, blue line). This reduced the mean absolute error (MAE) of the predicted PCr concentration from 3.28 mM to 1.05 mM and the pH value from 0.06 to 0.03. Notably, both estimated ATPase rates are below the theoretical maximal oxidative capacity of ATP synthesis in the model of 20 mM*/*min.

**Fig 3.**
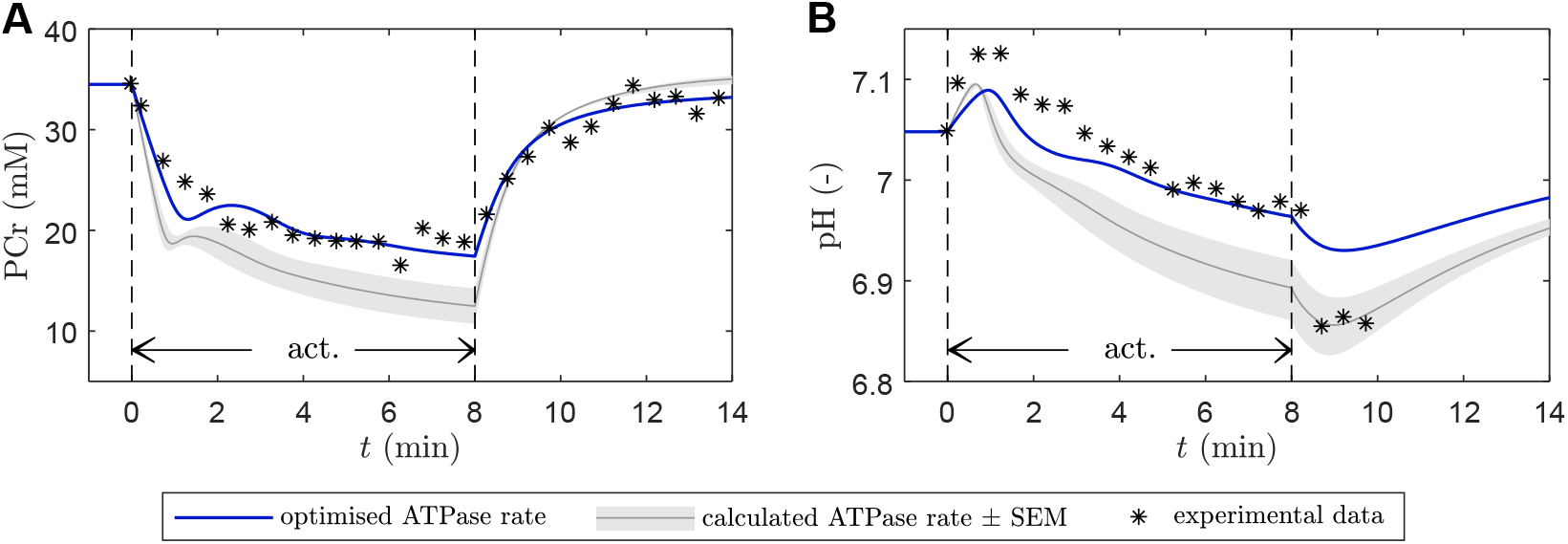
Model validation by comparing simulation results to independent experimental data [43].(A) [PCr] and (B) pH dynamics in response to a 8 min exercise protocol (electric stimulation with frequency 0.75 Hz). Black markers: experimental data [43]; shaded grey area: simulations with the range of ATPase rates given in [43]; blue lines: simulations with optimized ATPase rate. Note that the visualized PCr data is normalized such that the simulated and measured data have the same baseline.

#### 3.1.3 MPSA

To identify the parameters that have the largest influence on the model output and to test the robustness of the model given small perturbations of the parameters, we performed a local MPSA (see Section 2.2.4). Fig 4 summarises the results obtained from the MPSA. Thereby, only parameters with sensitivity values *D*_*m,n*_ *≥* 0.1 (with 1 indicating strong sensitivities and 0 denoting completely insensitive parameters) for at least one state variable at one of the muscle activation levels are presented. In the resting state, all baseline concentrations excluding ATP are most sensitive to pH (*D*_*m,n*_ *≥* 0.4). The basal ATP concentration is most sensitive to the size of the total adenine nucleotide pool (TAN), i.e., *D*_*m,n*_ = 0.47. Considering only active conditions, all state variables are most sensitive to the basal pH value (*D*_*m,n*_ *≥* 0.36). Further, only modest sensitivities of the model dynamics during muscle contraction are observed for variations of the phosphate (*D*_*m,n*_ *≤* 0.23), creatine (*D*_*m,n*_ *≤* 0.16), and adenine pool size (*D*_*m,n*_ *≤* 0.11), respectively.

**Fig 4.**
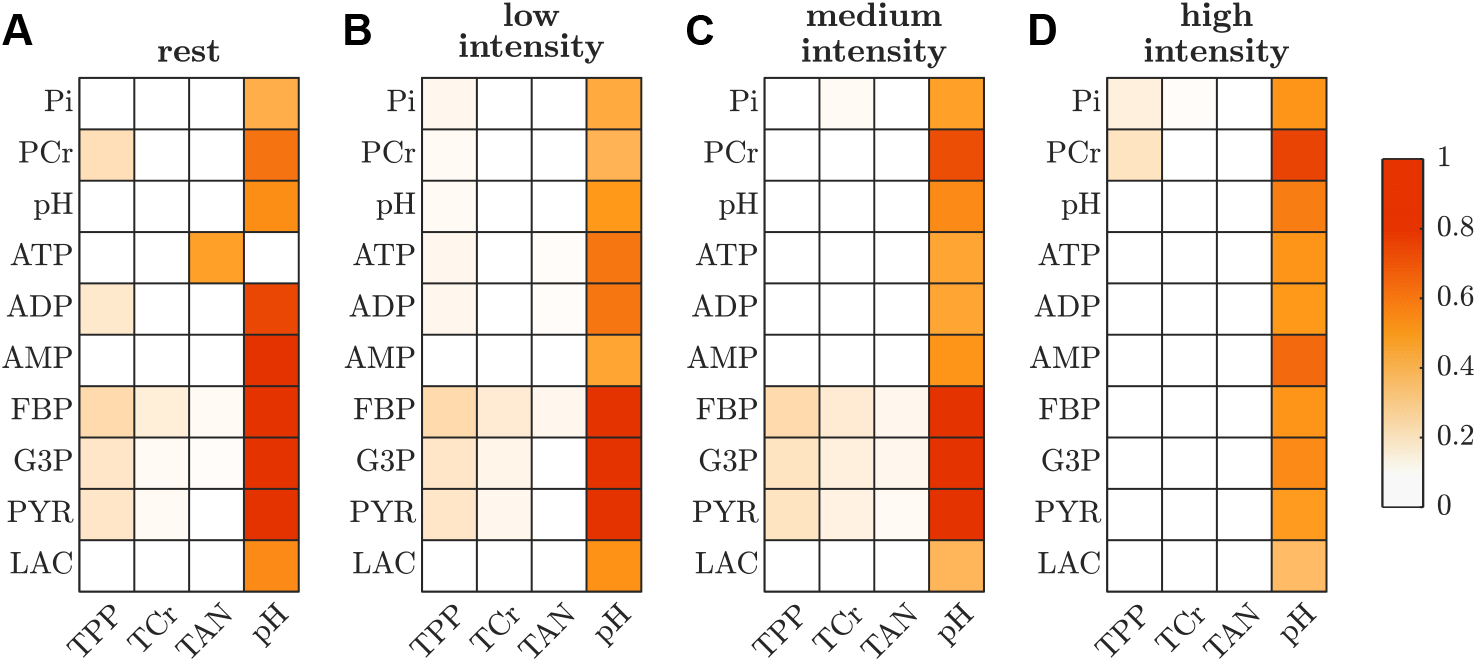
Summary of the multi-parameter sensitivity analysis (MPSA). The color scale indicates the Kolmogorov-Smirnov (K-S) statistic (*D*_*m,n*_ *∈* [0, 1]) between pairs of state variables *m* and parameters *n*, whereby higher values indicate stronger sensitivities. Columns represent the tested parameters with *D*_*m,n*_ *≥* 0.1 with respect to at least one of the state variables at any one of the simulated work rates. Rows show selected state variables. (A) sensitivity of the basal state. (B-D) sensitivity during muscle activations of variable intensity.

### 3.2 In silico studies

#### 3.2.1 Experiment 1: cellular redox potential

##### Temporal dynamics

The in silico model was used to predict the dynamics of the internal state variables at different contraction intensities (from 2.5 % to 150 % of the maximal oxidative ATP production capacity). Fig. 5 exemplarily illustrates the model response for a low, medium, and high-intensity contraction. This corresponds to ATPase rates below (50 %), right at (100 %), and above (150 %) the maximal oxidative ATP synthesis rate (20 mM*/*min). While ATP content is stable (approx. 10.3 mM) during all exercise protocols (Fig. 5B), PCr and pH decrease and free Pi increases considerably (Fig. 5A and Fig. 5C). In detail, the relative PCr concentration (with respect to the baseline PCr level) drops by approximately 49 %, 72 % and 93 %, while Pi shows an opposite trend with a 6-fold, 9-fold, and 11-fold increase in Pi content during the low, medium, and high-intensity work rates. Further, the pH values decrease to 7.0, 6.8 and 6.5 during low-intesnity, medium-intensity and high-intensity exercise, respectively. A transient alkalinization is observed at the onset of exercise, independent of the intensity of the exercise protocol.

**Fig 5.**
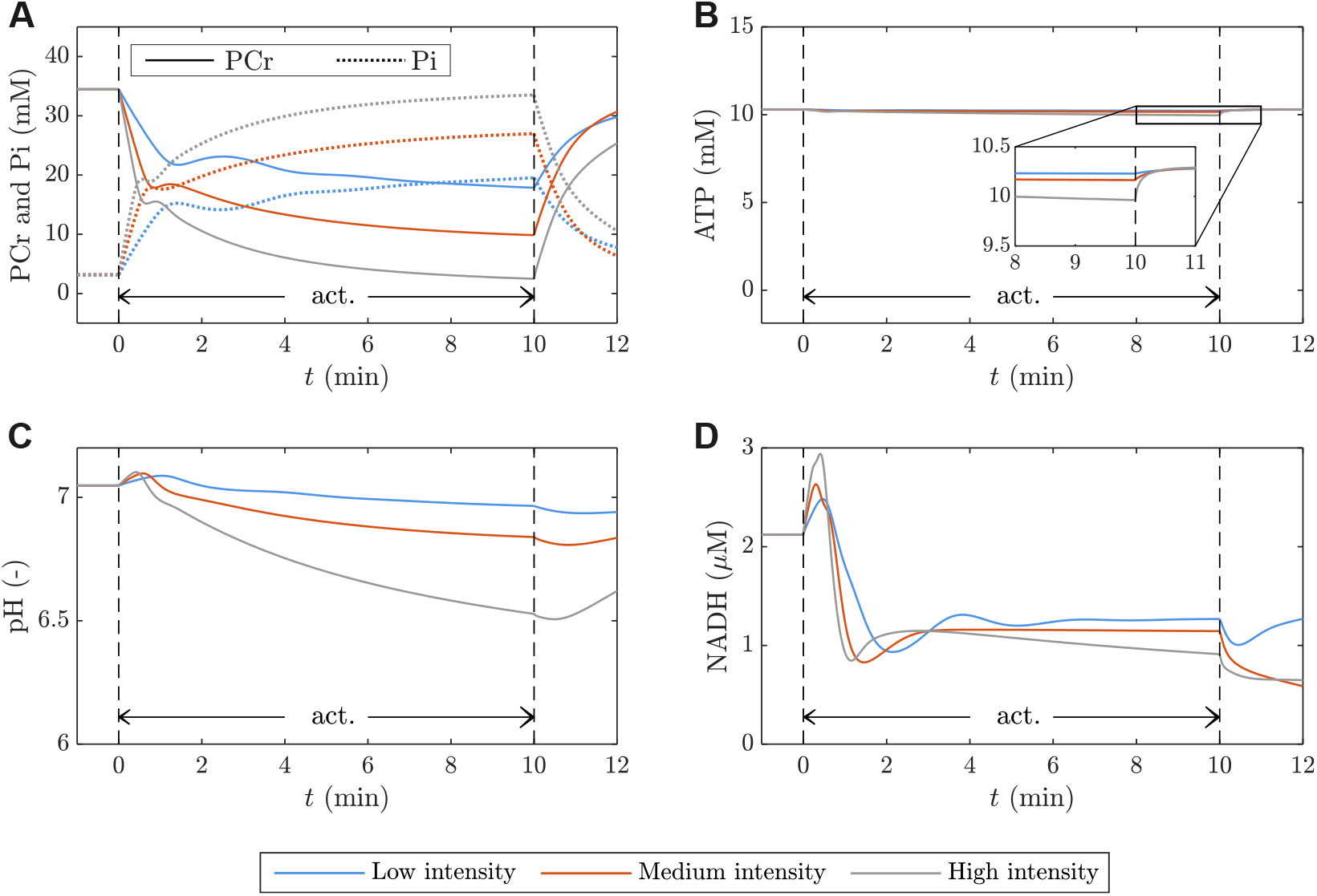
Model dynamics for three different contraction intensities (Blue: ATPase rate 10 mM*/*min, Orange: ATPase rate 20 mM*/*min, Gray: ATPase rate 30 mM*/*min). We show the measurable variables [PCr] (A), [Pi] (B), and pH (C) together with the non-measurable myoplasmic [NADH] (D).

The dynamics of the intracellular NADH concentration are presented in Fig. 5D. Directly after the onset of muscle contraction, a transient increase in the NADH concentration is observed. This is due to a slight imbalance between NAD reduction in the GAPDH and LDH (small reverse flux) reactions and the NADH oxidation through primarily OxPhos but also G3PDH (Supplementary Fig. 3A-C) resulting in a total net NADH generation flux in the micro Molar per minute range (Supplementary Fig. 3D-F). After a few seconds of muscle activation, an increase in the metabolic flux through GAPDH and subsequently through LDH, as well as activation of OxPhos, is observed. This leads to NADH re-oxidation such that the concentration drops below the basal level, i.e., approximately 50 % of the initial NADH concentration for all three work rates. Thereby, the NAD reduction via GAPDH is partially buffered by G3PDH. After a transient phase (i.e., approximately the first two minutes of muscle activation), the model virtually reaches a steady state.

##### Steady state

For low-intensity exercise the steady-state (*t* = 10 min) NAD^+^ resynthesis is approximately equally distributed between OxPhos and LDH (Fig. 6C). For medium and high-intensity contractions, the LDH contribution dominates. Notably, the oxidative pyruvate and NADH flux during low-intensity exercise (i.e., 50 % of the maximal oxidative capacity) is less than half of the maximum capacity of OxPhos for consuming NADH and pyruvate (i.e., 1.1 mM*/*min, whereby one makes use of the fact that the stochiometry factor of NADH and pyruvate is one, see Eqn. (14)). During high-intensity exercise, this value increases to around 80 % of the OxPhos reaction’s maximal flux rate.

**Fig 6.**
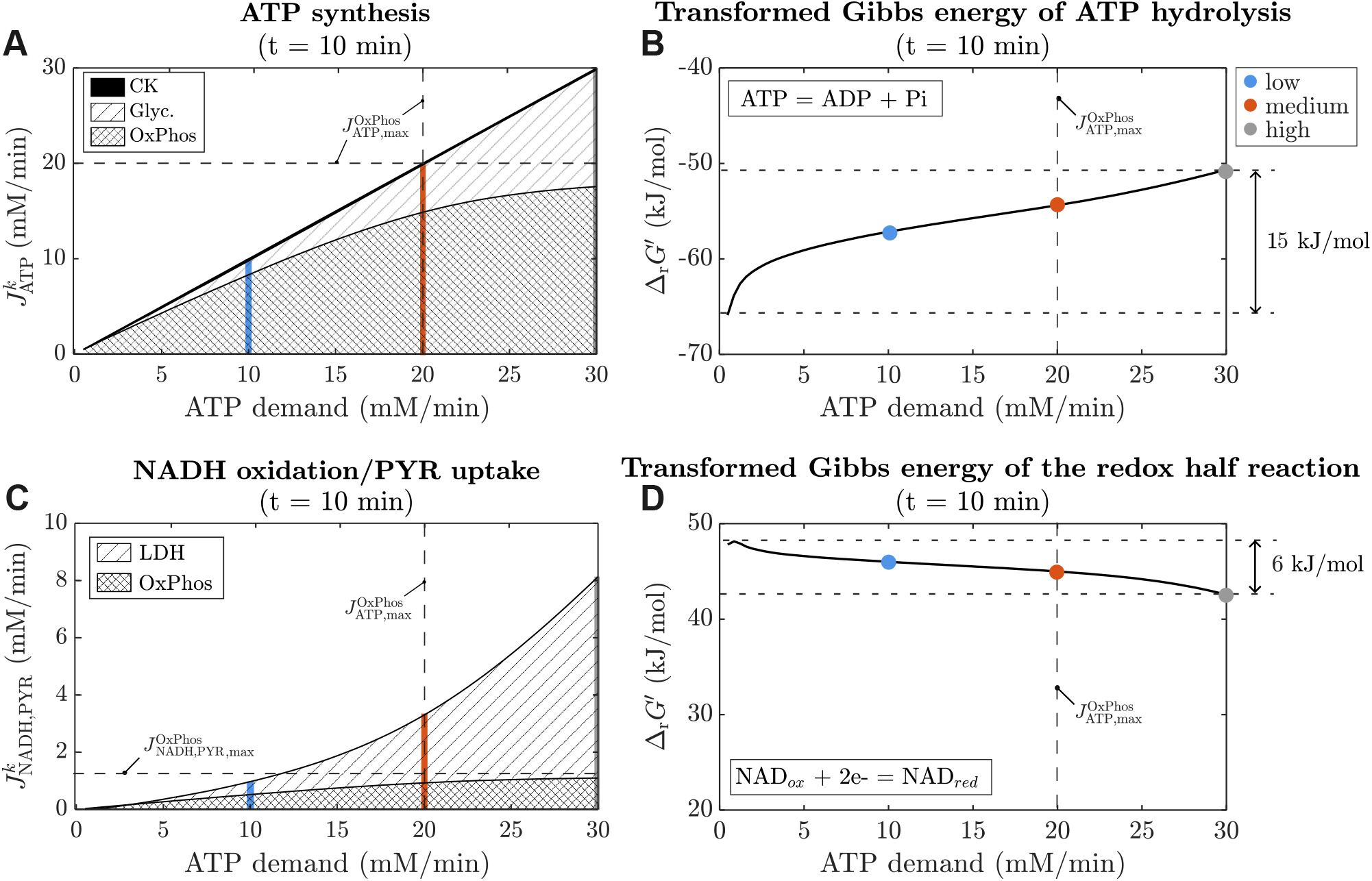
Contribution to (A) ATP synthesis and (C) NADH oxidation/PYR uptake as well as transformed Gibbs energy of (B) ATP hydrolysis and (D) the redox half reaction at the end of 10 min exercise at ATPase rates ranging from 0.48 mM (rest) to 30 mM. Colors indicate differnet ATPase rate: Blue: 10 mM*/*min, Orange: 20 mM*/*min, Gray: 30 mM*/*min.

Fig. 6A shows that for all contraction intensities, the ATP re-synthesis is dominantly achieved through OxPhos (low-intensity: 83 %, medium-intensity: 74 %, high-intensity: 59 %). Nevertheless, for the low-intensity and medium-intensity exercise below and at the maximal oxidative ATP synthesis capacity, the glycolytic pathway contributes considerably to ATP synthesis. Lastly, Fig. 6B and D show the transformed Gibbs reaction energy of the ATPase and NAD^+^/NADH redox (half) reaction (see Eqn. (11)), depending on the contraction intensity. Since the redox half reaction is not a stand-alone reaction in the model, 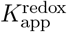 was calculated manually as described in Appendix D.3). It is observed that the change of the transformed Gibbs energy of ATP hydrolysis is nearly 3 times larger than the change of the Gibbs energy of the redox couple NAD^+^/NADH.

#### 3.2.2 Experiment 2: LDH knock-out

##### Temporal dynamics

Fig. 7 exemplarily compares the behaviour of the control model and the LDH-KO model given a 20-minute-long exercise protocol with an ATPase rate at the maximal oxidative capacity of ATP production 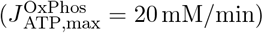. Both model configurations show stable ATP concentrations during the entire exercise protocol (Fig. 7A). The

**Fig 7.**
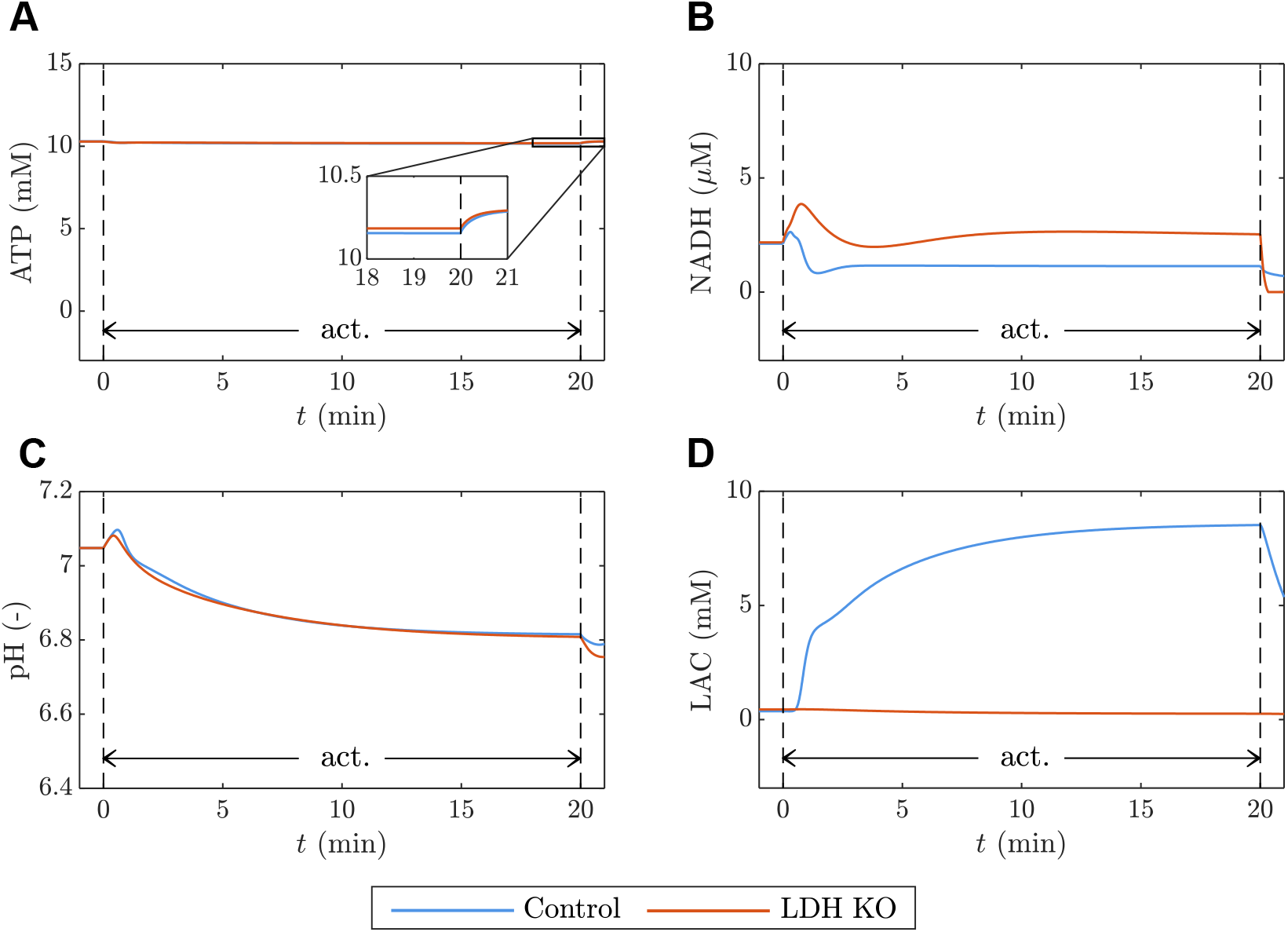
Comparison of the control model (blue) and LDH KO model (orange). (A) [ATP], (B) [NADH], (C) pH and (D) [LAC] dynamics during a 20 min exercise at an ATPase rate of 20 mM*/*min.

LDH-KO simulation lacks accumulation of lactate, whereas the intracellular lactate content in the control simulation increases to approximately 8.5 mM (Fig. 7D). Further, both the control model and the LDH-KO model show a transient reduction of the cytoplasmic redox state at the onset of muscle activation (Fig. 7B). The LDH-KO model shows a higher maximum NADH concentration (LDH-KO: 3.8 µM; Control model: 2.6 µM) and requires more time to reach a steady-state NADH concentration during exercise. Moreover, in the control model, the steady-state NADH concentration during exercise is below the baseline concentration (Δ[NADH] ≈ −1.0 µM), for the LDH-KO model, the steady-state NADH concentration is above the basal level (Δ[NADH] ≈ 0.4 µM). Lastly, we note that the difference in the predicted pH dynamics is marginal (Fig. 7C), with both systems dropping to a pH value of 6.8 (MAE = 0.006).

##### Steady state

We simulated 20 min long contractions with variable intensities (up to the maximum oxidative capacity of ATP synthesis, 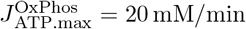) to predict the steadystate contributions of OxPhos and glycolysis to the overall ATP synthesis rate during exercise. The steady-state values were determined at the end of muscle activation (*t* = 20 min) and are summarized in Figs. 8B and 8E). For the control model and at the lowest considered ATPase rate (0.5 mM*/*min), OxPhos covers 99 % of the ATP demand. This value drops to 84 % and 75 % for ATPase rates of 10 mM*/*min and 20 mM*/*min, respectively. In the LDH-KO model, around 91 % of ATP demand is covered through OxPhos for all simulated ATPase rates.

**Fig 8.**
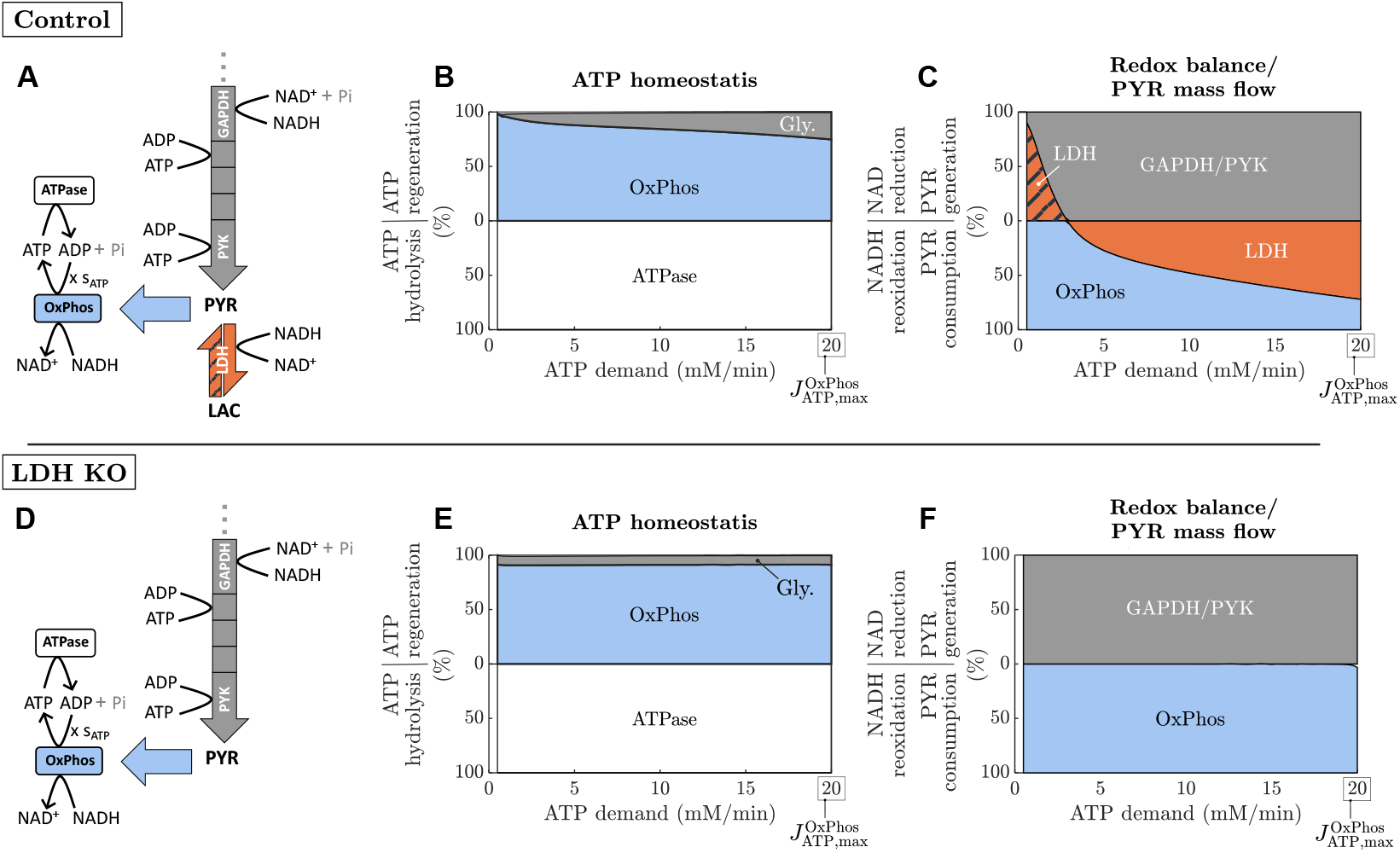
Comparison of (A) control model and (D) LDH KO model. Steady-state relative contribution to (B,E) ATP synthesis and (C,F) redox balance/pyruvate mass flow at the end of a 20 min exercise at ATP demands ranging from 0.5 mM*/*min to the theoretical maximal mitochondrial ATP synthesis rate of 20 mM*/*min.

Figs. 8C and 8F show the relative steady-state contribution of GAPDH, LDH, and OxPhos to cytoplasmic redox balance during exercise. In the control system and at the lowest considered ATPase rate (0.5 mM*/*min), LDH runs backward, i.e., utilizing extracellular lactate to generate pyruvate (though the flux is close to zero with −0.03 mM*/*min). Thereby, 91 % of total NAD reduction can be attributed to LDH. In contrast, for ATP demands above 3 mM*/*min, LDH converts pyruvate into lactate and, thus, together with OxPhos, oxidizes NADH. The relative contribution of LDH to NADH oxidation is around 48 % (LDH flux: 0.49 mM*/*min) and 72 % (LDH flux: 2.4 mM*/*min) for ATP demands of 50 % and 100 % of the oxidative ATP production capacity, respectively. Consequently, the metabolic system utilizes anaerobic glycolysis at ATPase rates considerably below the maximal oxidative ATP synthesis capacity. In the LDH-KO model 100 % of NADH generated by the GAPDH reaction is reoxidised via OxPhos.

#### 3.2.3 Experiment 3: LDH knock out under ischemia

To further study how myoplasmic redox balance impacts metabolite fluxes through the glycolytic pathway, we simulated one minute ATP metabolism at an ATPase rate of 20 mM*/*min under ischaemic conditions comparing the control model vs. the LDH-KO model. Fig. 9 (A-C) shows the averaged fluxes through each enzyme-catalysed reaction in the glycolytic pathway during muscle activation. Comparing the simulations of the LDH-KO model (Fig. 9B) with the control model (Fig. 9A), the glycolytic fluxes are elevated, particularly in the proximal part. For example, the flux through PFK increases by 422 %, while the reaction rate through PYK is increased by 55 %. Further, the LDH-KO simulation shows an increased activity of G3PDH by 711 % compared to the control system. The model predicts an 11-fold increase of the intracellular NADH concentration (Fig. 9E) as well as an accumulation of glycolytic intermediates (Fig. 9F) for the LDH-KO system during exercise. Compared to the control model there is no lactate accumulation, the pyruvate concentration at the end of muscle activation is 8.0 mM higher, the fructose 1,6-phosphate concentration increases by 9.9 mM and the glycerol 3-phosphate concentration is 7.5 mM higher. The altered dynamics in the LDH-KO model cause a negative net production of ATP in the glycolytic pathway, as the ATP consumption in the proximal part exceeds the ATP production in the distal part. That is, the model predicts a glycolytic ATP synthesis rate of −17.3 mM*/*min at the end of muscle activation compared to 17.0 mM*/*min in the control system. As a result, the system is not able to match the energy demand, leading to a 2 mM decrease in ATP (Fig. 9D).

**Fig 9.**
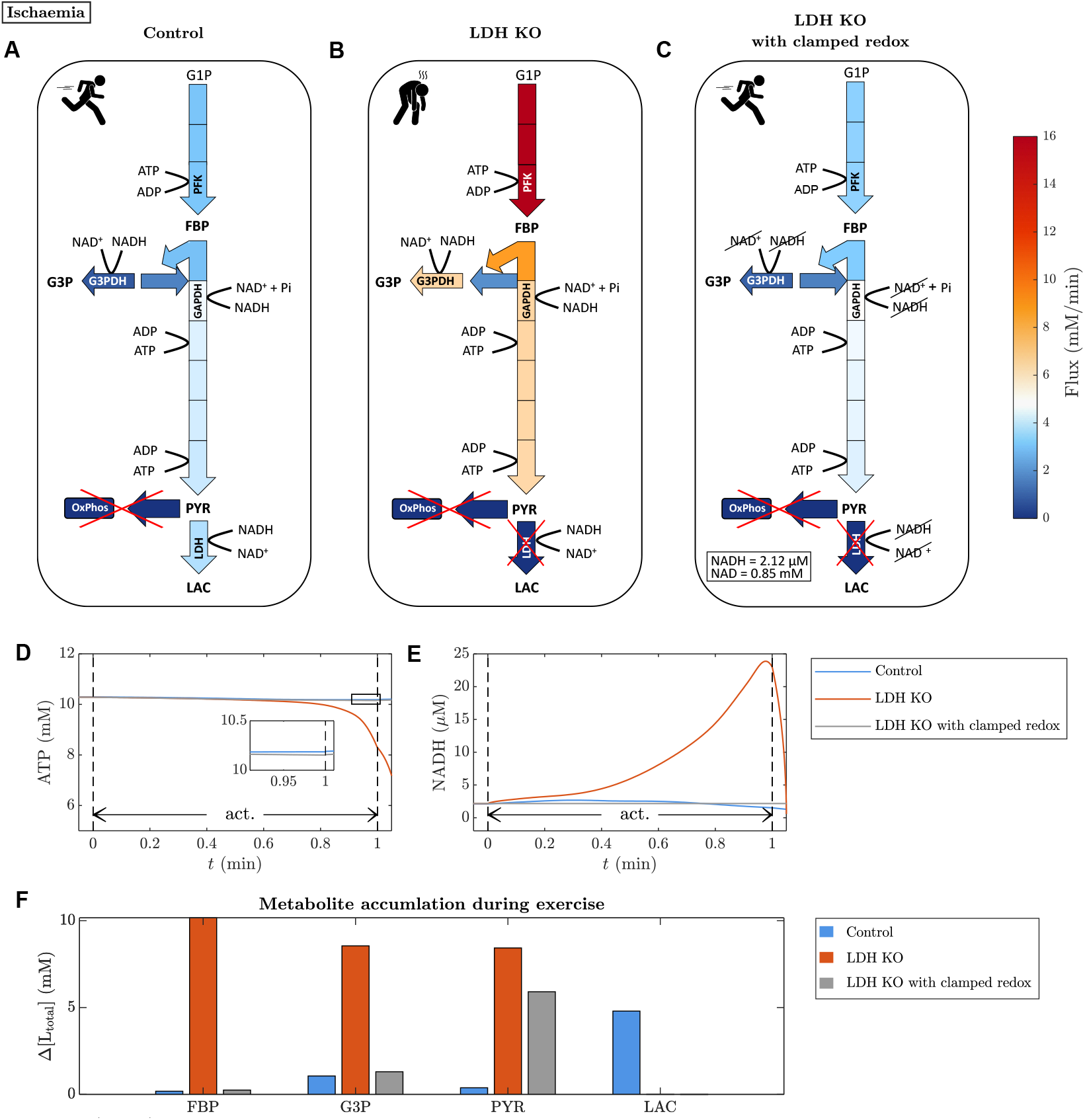
(A-C) mean enzymatic fluxes in the glycolytic reaction network in a simulation of one minute ischemic muscle exercise for three model configurations (A) control; (B) LDH knockout; (C) LDH knockout with clamped [NAD] and [NADH] (mean fluxes averaged over one min of simulation; ATPase rate = 20 mM*/*min; OxPhos rate = 0 mM*/*min). The magnitude of each individual enzymatic flux is colour-coded between 0 (dark blue) and 16 mM/min (dark red). (D) [ATP] dynamics and (E) [NADH] dynamics during one-minute ischemic muscle exercise. (F) Accumulation (Δ [L_total_] = [L_total_] (*t* = 1 min) − [L_total_] (*t* = 0)) of fructose-bi-phosphate (FBP), glucose-3-phosphate (G3P), pyruvate (PYR), and lactate (LAC) during one-minute ischemic muscle exercise.

To test the role of myoplasmic redox balance in maintaining sustainable glycolytic ATP production, we simulated an LDH-KO model with clamped NAD^+^ and NADH concentrations (Fig. 9C). This model configuration (LDH-KO, clamped redox) shows highly similar fluxes to the control model. For example, the PFK flux is elevated by 11 % and PYK reaction rate increases by 8 %. In particular, the activity of G3PDH only shows a moderate increase of 24 %. Accordingly, compared to the control model, there is no considerable accumulation of glycolytic intermediates (Fig. 9F). The most relevant difference is the increased level of pyruvate and the lack of lactate accumulation, both concomitant effects of the LDH knock-out. Analogous to the control model, the model with clamped NAD^+^ and NADH is able to maintain ATP homeostasis (Fig. 9D) with a positive net production of ATP in the glycolytic pathway. At the end of the one minute long exercise, the glycolytic ATP synthesis rate is 11.9 mM*/*min.

## 4 Discussion

This study identifies and validates a computational model of carbohydrate metabolism in active FOG myofibers based on dynamic in vivo ^31^P-MRS recordings of ATP metabolite and pH time courses. The model provides novel insights into the concomitant dynamics of metabolic intermediates, including pyruvate and myoplasmic NADH, that are difficult to measure experimentally. The presented simulations make novel predictions on the role of pyruvate and lactate metabolism in skeletal energetics. For example, analysis predicts that the contribution of lactate dehydrogenase (LDH) to NADH redox balance is crucially important in maintaining ATP homeostasis when metabolic demand exceeds the oxidative ATP synthesis capacity. In fact, we predict that LDH’s function in maintaining relatively low cytoplasmic NADH levels is fundamentally more important than its role in slowing cellular acidification under high ATP demand. Moreover, the presented simulations reveal that a purely feedback-driven biochemical control scheme is able to explain the ATP supply-demand relationship over a roughly 100-fold range of ATP demands.

### 4.1 ATP turnover-driven carbohydrate metabolism in the FOG myofiber phenotype

Computer simulations of a FOG myofiber’s response to step changes in myoplasmic ATP demand show that the combined action of carbon mass flow and feedback control of OxPhos embedded in the model suffices to maintain stable ATP and energetic state over a 100-fold range of ATP turnover rates (Fig. 2). The predicted relative contributions of glycolytic and oxidative ATP synthesis flux to total ATP production over ATP turnover rates spanning from basal rate to 1.5-fold above the maximal rate of mitochondrial OxPhos (Fig. 6A) are in line with estimates from experimental in vivo studies [48–50]. Likewise, the predicted magnitude of changes in myoplasmic PCr and Pi concentrations and pH with increasing ATP turnover (Fig. 5) associated with a 15 kJ/mole drop in free energy of myoplasmic ATP hydrolysis over this operational range (Fig. 6) are similar to in vivo experimental observations in skeletal muscles composed predominantly of FT myofibers [51–54]. As such, these results support the contention of previous in silico studies of energy balance in skeletal muscle that feedback control of mitochondrial OxPhos suffices as a coarse regulatory mechanism to maintain ATP energy balance in this tissue [55, 56]. Earlier studies that proposed that some form of feedforward control of mitochondrial ATP synthesis must additionally be operational in vivo to balance high rates of cellular ATP turnover [39, 57] only considered feedback control by ADP. It has since been shown that Pi contributes to respiratory control in FT myofibers [58, 59] and was included in the present model.

These model simulations inform the understanding of the phenomenon of ‘aerobic lactate production’ in skeletal muscle [5]. This term refers to the energetically seemingly wasteful conversion of pyruvate to lactate despite presumed adequate oxygen availability and mitochondrial OxPhos capacity first observed in cancer cells [60] and later in other cell types, including skeletal myofibers (e.g. [61, 62]). A recent study in cancer cells concluded that insufficient capacity of mitochondrial redox shuttles to buffer elevated glycolytic NADH production drives lactate production in this particular glycolytic cell type and proposed this outcome may be generalizable to other cell types, including skeletal myofibers [9]. Our findings reject the latter contention: LDH knockout simulations show that oxidative recycling of myoplasmic NADH in the FOG myofiber phenotype in and by itself suffices to maintain redox balance over the aerobic range of ATP turnover rates (Fig. 8), accompanied by faster and steeper on-kinetics of OxPhos (Supplementary Fig. 5A-B). This outcome is in agreement with empirical findings that patients with muscular LDH deficiency exercising at low to moderate intensities experience no symptoms [63, 64]. Simulations of the intact metabolic model reveal aerobic lactate production in this myofiber phenotype is driven by the mass action effect of LDH substrate accumulation shortly after activation of glycolysis by ATP turnover (Supplementary Fig. 4). Indeed, the majority of lactate produced at low and intermediate ATP turnover rates was predicted to occur in this early time frame of transition to a new steady state (see Supplementary Fig. 3). Further, the predicted behaviour of the LDH-KO model under ischemia (Fig. 9) is in agreement with studies on LDH deficiency, where patients experience muscle cramping and myoglobinuria (dominantly) induced by ischemic exercise due to an inability to maintain ATP homeostasis [63, 64]. Muscle biopsies revealed increased concentrations of pyruvate and glycolytic intermediates (e.g., fructose 1,6-biphosphate (FBP)) after ischemic exercise while lacking the characteristic lactate accumulation. In particular, observed higher levels of glycerol 3-phosphate (G3P) were attributed to an increased activity of G3PDH. Clamping the myoplasmic redox state, the computer simulations indicate that intolerance of high-intensity exercise in LDH-deficient patients is mainly due to failing homeostasis of myoplasmic redox balance rather than accumulation of glycolytic intermediates in the distal part of the pathway.

Therefore, the picture that emerges from these simulations is that aerobic lactate production in FOG myofibers is simply a byproduct of metabolic flexibility associated with expressing high levels of LDH, affording continual ATP synthesis under conditions of compromised vascular oxygen supply or high intensity exercise. A similar view has previously been proposed by others (e.g. [6]) albeit using qualitative reasoning on the basis of the high (i.e., 10^4^) value of the apparent equilibrium constant of the LDH reaction. The LDH reaction is crucial to maintaining cytoplasmic redox balance under conditions where metabolic demand exceeds oxidative ATP synthesis capacity. Note that any lactate produced by FOG myofibers is not lost to the body but made available as oxidative substrate to other cells and tissues including neighboring red myofibers (e.g. [65,66]). Thus, the phenomenon of aerobic lactate production does not necessarily represent a metabolic inefficiency at the level of the whole organism. Moreover, our findings offer insights into two lingering debates in skeletal muscle biochemistry. First, regarding the longstanding debate on whether or not LDH conversion of pyruvate contributes to myofiber acidification (e.g., [67–69]), our LDH knockout simulations of myoplasmic pH dynamics (Fig. 7C) predict that this reaction in and by itself has little influence on cellular acid-base balance [70, 71]. Second, regarding contentions that lactate is always the product of glycolysis [6, 72], our model simulations indicate that the latter is untenable in absolute sense. Steady-state myoplasmic concentrations of pyruvate and lactate during exercise are both predicted to progressively increase with increasing ATP turnover (Supplementary Fig. 4). In fact, our model simulations predict that immediately upon a step change in ATP turnover rate, any lactate in the myoplasm or interstitial matrix is converted to pyruvate in order to fuel mitochondrial ATP production while glycolysis is ramping up (Supplementary Fig. 3). This particular model prediction may, however, alternatively reflect the absence of any feed-forward control of glycolytic flux in the present model. Two previous studies of glycolysis in skeletal muscle have proposed that sensing and transduction of the step change in myoplasmic calcium concentration upon myofiber recruitment that precedes any increase of ATP turnover rate may indeed be operational in vivo [19, 59].

A final main result of step response testing of our model is the finding of second-order underdamped dynamic behaviour of the network in transitions between steady states, manifest, for example, in the time course of myoplasmic concentrations of PCr, Pi and NADH (e.g. Fig. 5). This behaviour is observed at all tested ATP turnover rates, but most prominently at low ATP turnover rates. Its origin is a transient mismatch between carbon mass flow through glycolysis and mitochondrial OxPhos, respectively, resulting from ADP stimulation of both PFK and OxPhos (Appendix B.1.3 and main text Section 2.1.2). This interpretation is confirmed by the finding that increasing *V*_max_ of OxPhos or LDH knockout aggravated the second-order behaviour by increasing pyruvate mass flow to mitochondrial metabolism (Supplementary Figs. 7 and 5) while decreasing mass flow through lower glycolysis. Increasing *V*_max_ of G3PDH had the opposite effect (Supplementary Fig. 7). Comparison of model simulations of PCr and Pi dynamics to experimental in vivo ^31^P-MRS recordings from FT muscle at low to moderate electrical stimulation frequencies tend to support the model prediction of second-order behaviour (Fig. 2A-B and D-E) but this interpretation is inconclusive due to signal-to-noise limitations inherent to dynamic in vivo ^31^P-MRS [73]. Alternatively, this particular model prediction may be construed as demonstrating that kinetic controls of PFK and other glycolytic enzymes not captured by our model in fact play a role in fine-tuning carbon mass flow through upper and lower glycolysis in vivo. Specifically, our model does not capture the complete set of known allosteric modifiers of the enzymatic activity of PFK and GAPDH [74] nor any of the proposed interactions of these enzymes with cytoskeletal filament proteins or calcium-binding proteins [75, 76] that have been proposed to modify glycolytic activity in vivo [77]. As such, the present model prediction of underdamped second-order behaviour of the metabolic network for G1P metabolism in an FOG myofiber in response to a step change in ATP turnover rate awaits experimental verification.

### 4.2 Modeling muscle energetics

Computational modeling has a rich tradition in muscle energetics. Here, we discuss the proposed model in the context of the existing literature and potential applications.

#### 4.2.1 Model improvements and comparison to existing models

Similar to previous models of ATP metabolism in muscle fibers (e.g., [39, 56, 78]), the proposed model incorporates the key components of ATP homeostasis - i.e., ATP turnover by cytoplasmic ATPases and ATP synthesis by creatine kinase and oxidative phosphorylation, respectively, linked by a feedback control loop. In addition, the proposed modelling framework incorporates a detailed model of G1P catabolism including redox coupling to OxPhos, adenylate kinase buffering of ATP/ADP, LDH buffering of cytoplasmic redox, lactate exchange with extracellular space and computation of the emergent state variable cytoplasmic pH building on previous work by Vinnakota and coworkers in their study of resting biochemical state in an inactive muscle fiber [12]. Thus, the proposed modelling framework sets itself apart from existing computational models of energy metabolism in skeletal muscle that only focus on individual pathways (e.g. [12, 55, 74, 77, 79, 80]) as well as previous modelling efforts of coupled glycolytic and oxidative ATP synthesis that are limited to the resting state [81] or dismiss thermodynamic constraints and/or the effects of pH dynamics (e.g. [82–84]). A detailed model of coupled glycolytic and mitochondrial energy metabolism in skeletal muscle with a mixed feed-forward and feedback control structure of high complexity was previously developed to investigate dynamic behaviour during exercise. However, only comparatively small jumps in ATP demand were considered, and the validity of the dynamic model response was tested against steady-state values [85].

### 4.3 Limitations

The presented results are based on a mathematical model capturing the pH dependency and thermodynamics of the simulated biochemical reactions. Nevertheless, the presented model is a simplification of the underlying physiological system. The associated limitations are discussed in the following.

#### 4.3.1 Uncertainty of parameters and validation

The proposed model has a high number of variable model parameters with N = 85 (listed in Supplementary Tables 2 and 3). Yet, a particular strength of the modelling framework is that most parameters can be fixed based on data from in vitro experiments (i.e., a data-rich class of data) together with thermodynamic constraints (i.e., first principles). Nevertheless, there exist a few parameters for which appropriate experimental data is currently not available. These parameters were either obtained by utilizing values from measurement conditions slightly deviating from the simulated conditions (e.g., the CK equilibrium constant), fitting the model to ^31^P-MRS data (see Section 3.1.1) or heuristic parameter adjustments yielding consistency with the expected system behaviour (e.g., the PFK regulation). Particularly the latter yields parameters subject to considerable uncertainty. Further, the model parameters were obtained from different species (e.g., rat, human, rabbit, mouse, frog) and muscles (e.g., gastrocnemius, gracialis, flexor digitorum superficialis). Since enzyme activities vary between species and muscle (fiber) type [86], the model parametrisation was derived from fast twitch mammalian skeletal muscle, if available. We further note that the model showed the highest sensitivity to changes in the myoplasmic pH (Section 3.1.3). Most maximum enzyme activities were empirically corrected given the current pH value (cf. [12]), however, such data was not available for all considered reactions.

While the model predicted behaviours, closely replicating observation from ^31^P-MRS measurements, direct validation of the predicted internal states (e.g., cytoplasmic redox state and glycolytic intermediates) is currently not possible due to methodological limitations. Yet, the in silico experiments presented in Fig. 8B and Fig. 9 showcase that the model is able to qualitatively capture the experimentally observed behaviour, e.g., the effect of perturbations in redox balance caused by LDH deficiency.

#### 4.3.2 Kinetic models

All enzyme kinetics in the proposed model were adopted from the literature [15,17,23,79,81]. The majority of enzyme-catalysed reactions are modelled based on a rapid equilibrium assumption for on and off binding-reaction kinetics between metabolites and enzyme. This assumption might have limitations for flux rates close to the maximum enzyme activities. Further, enzyme-catalysed reactions are assumed to be reversible and thermodynamically constrained, satisfying the Haldane relation. Exceptions are the kinetic descriptions of ATPase, PFK and OxPhos. Their limitations are specifically discussed in the following.

##### ATPase model

The kinetic description of ATPase is a superposition of a fixed basal rate and an activity-dependent contribution. The simplification of a fixed activity-dependent ATP turnover misses regulatory mechanisms in response to fatiguing exercise. Thus, when simulating extreme scenarios, e.g., high-intensity exercise to exhaustion, the model can become numerically unstable. This can potentially be fixed by using a more detailed ATPase model.

##### Glycolytic substrates

We note that energy substrate supply through blood glucose and thus the hexokinase reaction (i.e., converting glucose to G6P) are not considered. Further, due to the lack of a suitable kinetic model, glycogen phosphorylase (GPa and GPb) is not explicitly modelled in the simulated reaction network. Thus, PFK is the first rate-limiting enzyme of the glycolytic pathway. This limits the reliability of the model predictions, particularly for the dynamics of G6P and F6P.

##### PFK model

Despite being intensely studied, there is still a lack of a thermodynamically constrained model of PFK kinetics considering the (coupled) regulatory effects of multiple metabolites such as ATP, ADP, AMP, Pi, and pH dynamics. The utilized kinetic model [15, 87] describes an irreversible biochemical reaction, i.e., without thermodynamic restriction. Under physiological conditions, PFK is considered to be irreversible (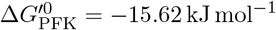 [88]). Yet, the lack of thermodynamic constraints may cause non-physical behaviours; specifically when simulating strong perturbations in ATP balance. Thus, we tested the thermodynamic consistency by calculating the transformed free energy of the PFK reaction. We observe that 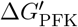 remained negative throughout the simulated period, verifying the thermodynamical consistency of the model within the boundary of the considered conditions.

##### Mitochondrial model

The kinetic description of the OxPhos model is based on an empirical sigmoid Hill function with feedback regulation through ADP [23]. We added regulation via inorganic phosphate and pyruvate and found that Pi regulation is necessary to prevent a complete depletion of inorganic phosphate during recovery. Yet, the utilized phenomenological model provides limited insights into the function of mitochondria. This would require a more detailed OxPhos model (e.g. [89]). While the proposed model describes a FOG fiber, another limitation of the implemented OxPhos model is the lack of fatty acid oxidation to fuel the resting state.

#### 4.3.3 Substrate exchange between compartments

The literature regarding the kinetics of carbon dioxide as well as lactate transport is sparse. Thus, their implementation is subject to considerable uncertainty. Further, we note that the considered transport mechanisms are a simplification of mass exchange in real muscle fibers, e.g., the selected MCT model discards the symport of non-lactate monocarboxylates, such as pyruvate. Moreover, the presented simulations neglect oxygen dynamics and its regulatory effects on the metabolic network. Compromised oxygen supply to myofibers during high-intensity work is, however, not uncommon [4] and oxygen availability is highly dynamic. For example, we do not consider vascular recruitment during muscle activation [90] and fiber-type specific capillarization [8]. Lastly, note that clamping the extracellular lactate concentration to 1 mM overestimates lactate removal from the extracellular space during high-intensity exercise (e.g. [91]).

### 4.4 Conclusion

Within this work we have developed a mathematical model for the quantitative investigation of energy metabolism in working fast-twitch oxidative glycolytic (FOG) fibers. Besides measurable metabolic variables, the model can predict the dynamics of variables that have proven challenging to probe experimentally in an intact physiological system, e.g., myoplasmic redox state and reaction fluxes. The presented simulations yield novel insights into how reported experimental observations such as aerobic lactate production in muscle emerge from the integrative behaviour of the metabolic network and are essential for maintaining robust energy and redox balance across a 100-fold working range. We anticipate that the proposed simulation environment will further enhance our understanding of muscle fatigue in health and disease and guide the development of treatment strategies of diverse neuromuscular disorders.

## A SUPPLEMENTARY MATERIALS

## B Mathematical model

Each flux is modelled as a function of metabolite concentrations associated with the corresponding biochemical reaction. If not disclosed otherwise, biochemical reactions are reversible and thermodynamically constrained by the Haldane relation as described in Lambeth and Kushmerick [79]. Exceptions are the kinetic equations of ATPase, PFK, and OxPhos.

We used the Biochemical Simulation Environment, BISEN (Medical College of Wisconsin, WI, USA) [36], to generate the system of differential equations. The effects of proton binding on the thermodynamics and kinetics of biochemical reactions are accounted for as proposed by Vinnakota et al. [92]. The hereto necessary reference reactions to each biochemical reaction are listed in Supplementary Table 1. The forward direction was defined as the reaction proceeding from left to right. The 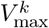 values reported in Table 1 (main text) were modelled as functions of pH using an empirical approach, for enzyme activities with known pH dependence over the physiological range. Thereby, the maximal enzyme activity at a specific 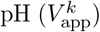 is the productof the maximal activities reported in Table 1 (main text) and an normalized algebraic function with its maximum at pH optimum [12, 81].

### B.1 Flux expressions

#### B.1.1 ATP hydrolysis

We modelled ATPase activity as an irreversible reaction, entering the model as a boundary flux. The reaction flux is the superposition of a fixed basal rate 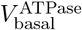 [13] and a step function with variable amplitude 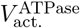 depending on the work rate (WR_*j*_) in a time interval *A*_*j*_ = { *t ∈* ℝ | *t*_*j*_ *≤ t < t*_*j*+1_}:

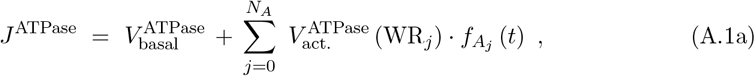

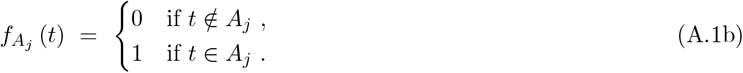

Therein, *N*_*A*_ *≥* 0 denotes the number of simulated exercise phases and 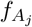 denotes the interval indicator function. 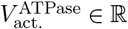 is the model input variable.

**Supplementary Fig. 1.**
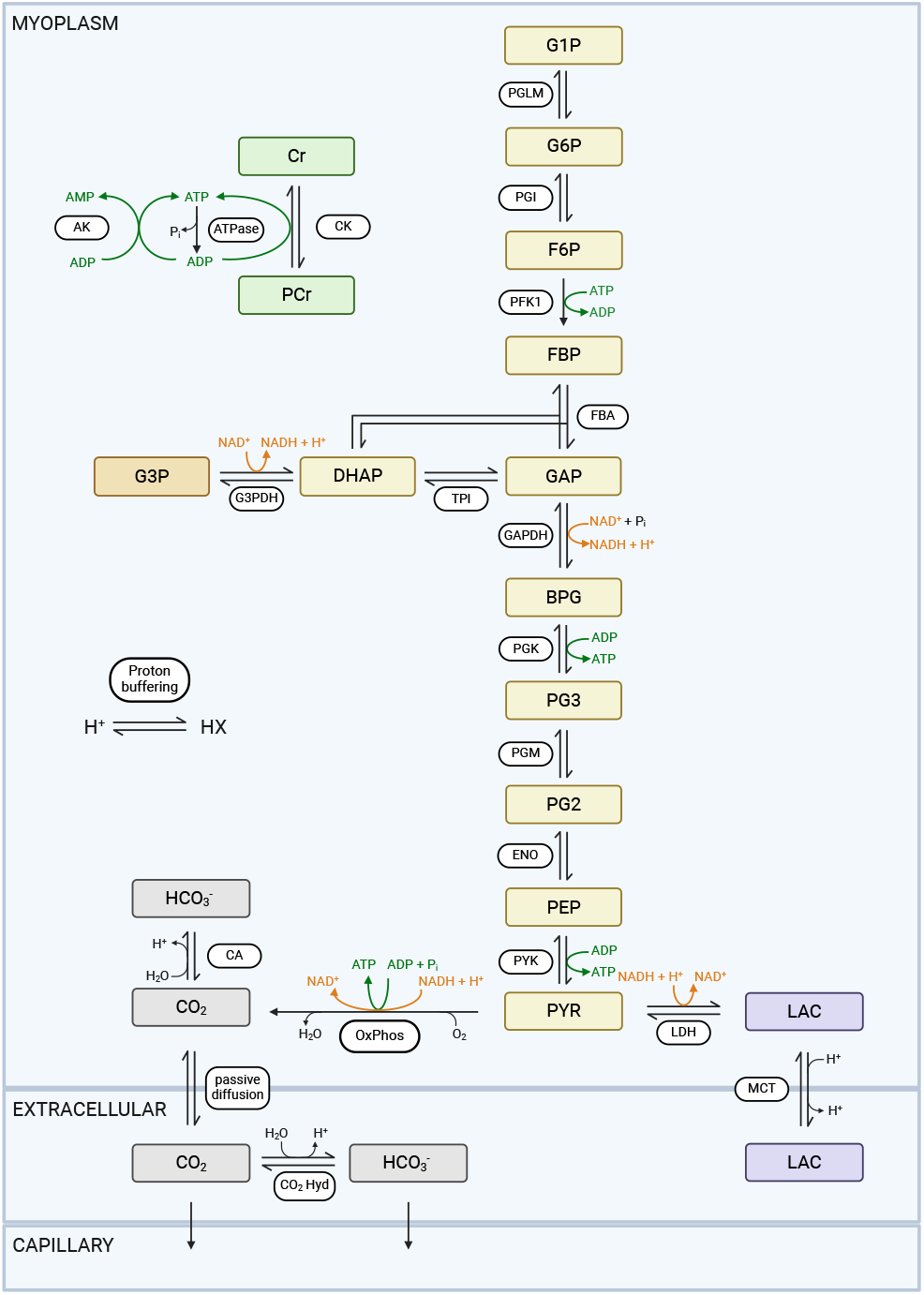
Schematic overview of the computational model. The model consists of a myoplasmic compartment (myoplasm), a compartment representing the interstitial fluid (extracellular) and a capillary compartment (capillary). The metabolic reaction network comprises of a lumped ATP hydrolysis rate (ATPase) balanced by ATP re-synthesis through (i.) the buffer reactions catalysed by creatine kinase (CK) and adenylate kinase (AK), (ii.) glyco(geno)lysis with every enzyme-catalysed reaction modelled individually starting from phosphogluco mutase (PGLM) to lactate dehydrogenase (LDH) and (iii.) a phenomenological description of oxidative phosphorylation (OxPhos). Additional redox buffering through the glycerol-3-phosphate dehydrogenase (G3PDH) reaction and proton buffering through the carbonic anhydrase (CA) reaction and an intrinsic cellular proton buffer are considered. The model also accounts for facilitated lactate transport via monocarboxylate transport (MCT4) and CO_2_ diffusion between the myoplasmic and extracellular compartment as well as CO_2_ and bicarbonate washout from the extracellular space into the capillary compartment. The figure was created using BioRender.com.

**Supplementary Table 1.**
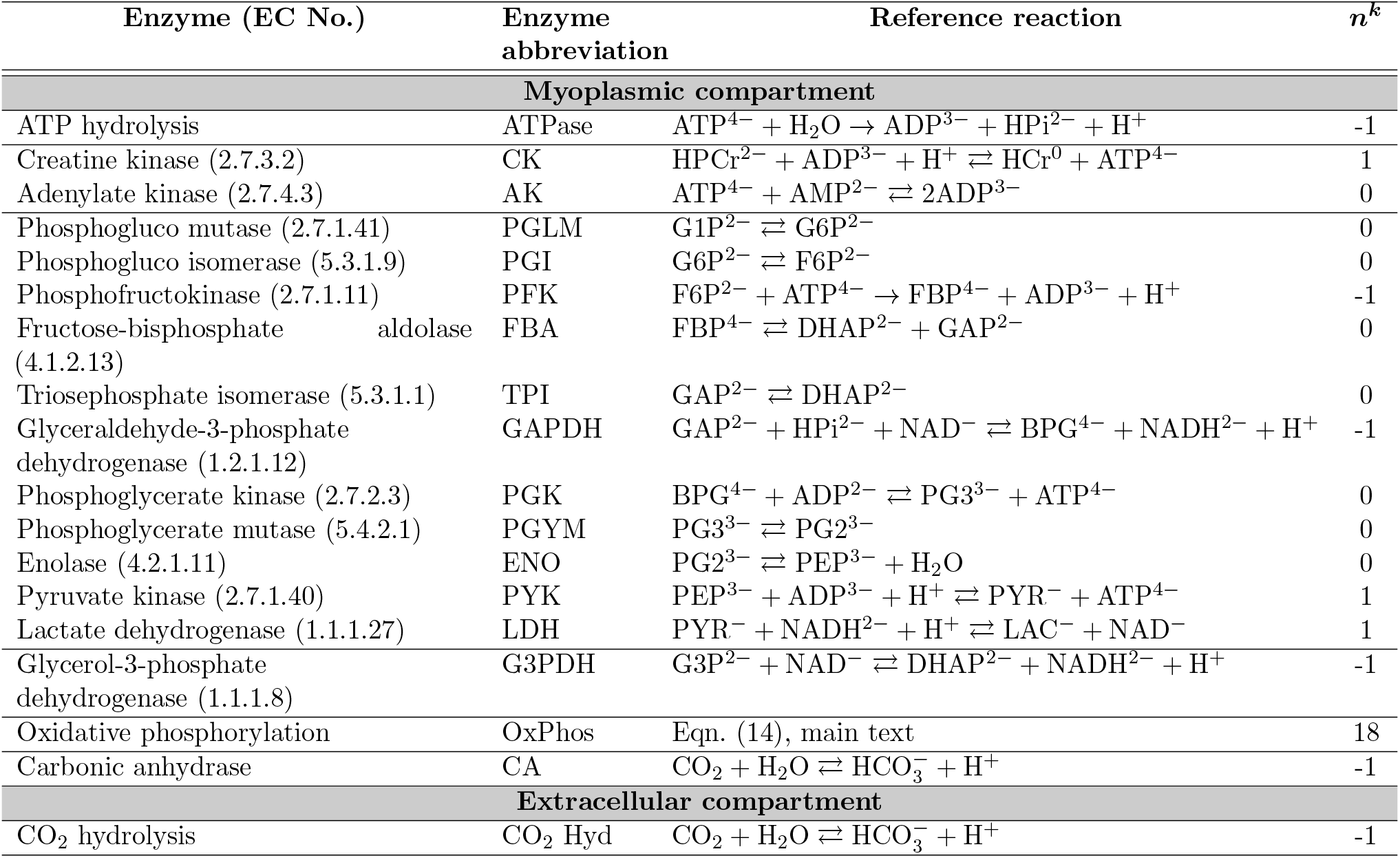
Reference reactions.

#### B.1.2 ATP buffer reactions

##### Creatine kinase, CK

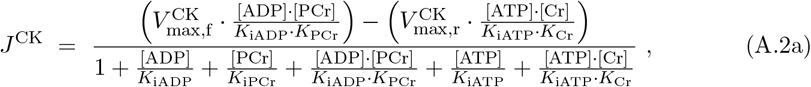

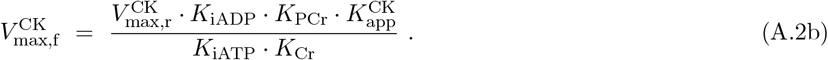

##### Adenylate kinase, AK

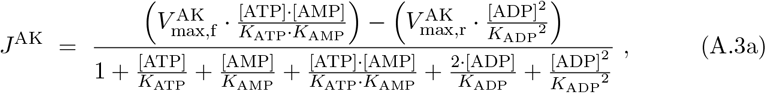

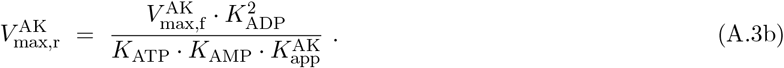

#### B.1.3 Glycolytic pathway

##### Phosphoglucomutase, PGLM

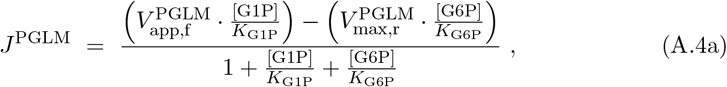

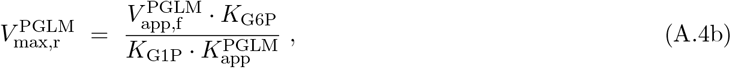

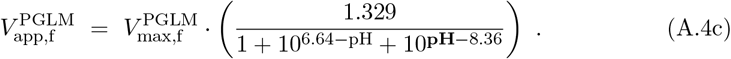

##### Phosphoglucoisomerase, PGI

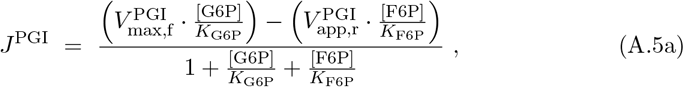

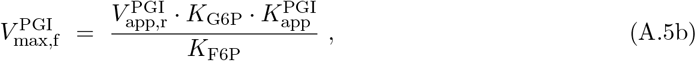

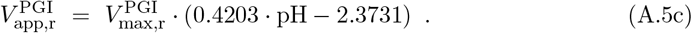

##### Phosphofructokinase, PFK

The kinetic model description for PFK originates from Waser et al. [15] and includes an adaptation to the 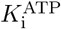 term by Connett [17] to account for competitive ADP and AMP activation to the ATP inhibition. The term was further adapted by dismissing a competitive activation via Pi, which was originally included in Connett’s adaptation, and considering the deinhibition effect of ADP instead of MgADP [77]. All parameter of the PFK model are provided in Supplementary Table 2. Note that 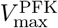 is not modelled as a function of pH.

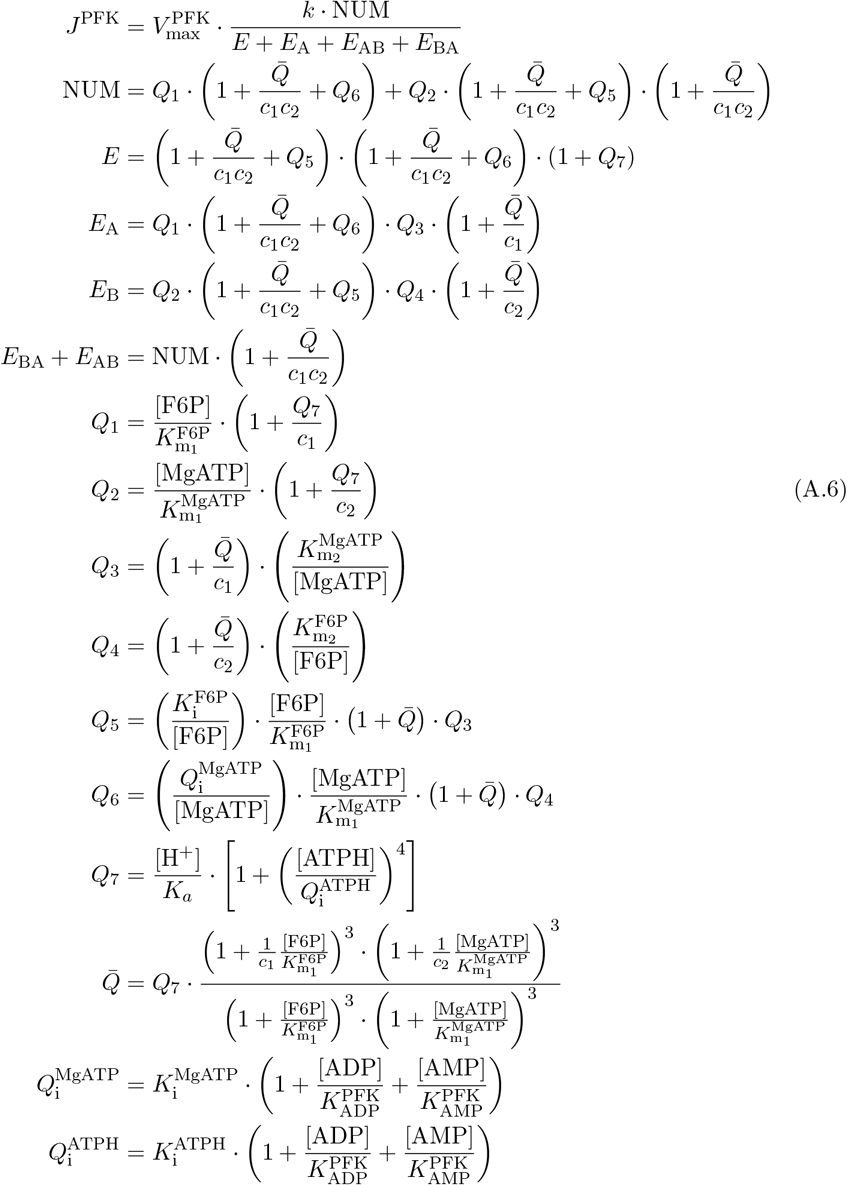

##### Fructose-bisphosphate aldolase, FBA

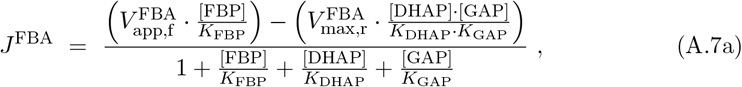

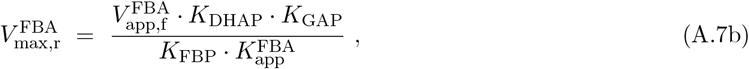

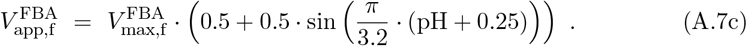

##### Triosephosphate isomerase, TPI

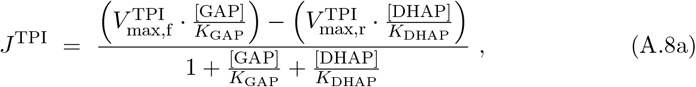

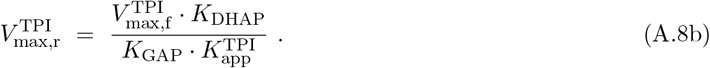

##### Glyceraldehyde-3-phosphate dehydrogenase, GAPDH

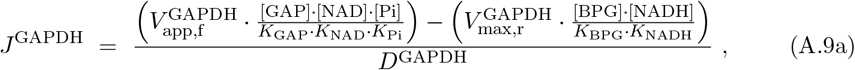

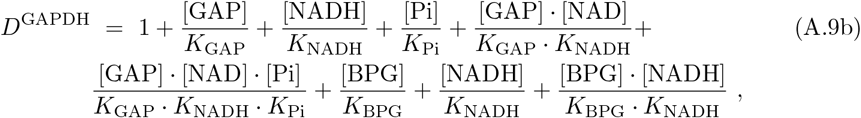

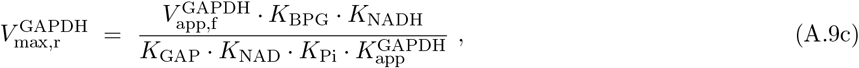

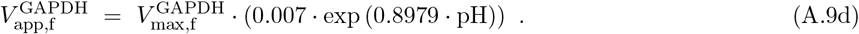

##### Phosphoglycerate kinase, PGK

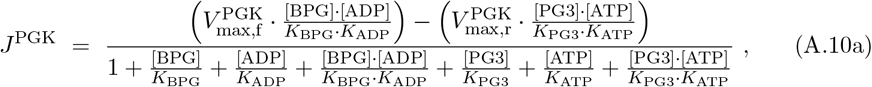

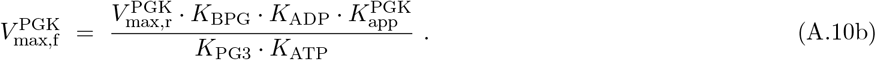

##### Phosphoglycerate mutase, PGYM

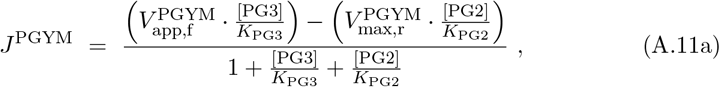

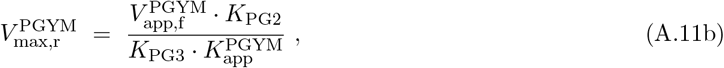

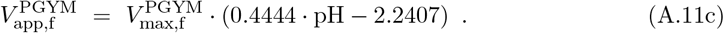

##### Enolase, ENO

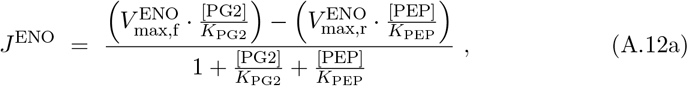

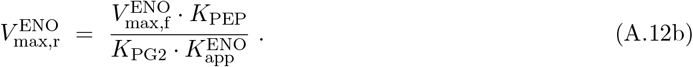

##### Pyruvate kinase, PYK

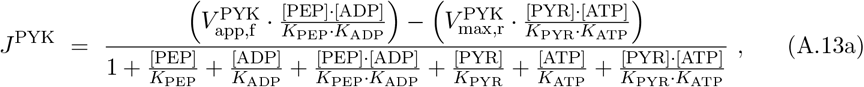

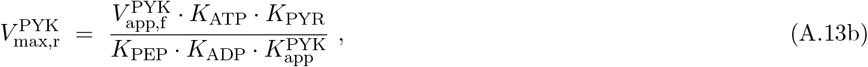

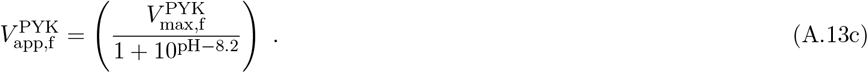

##### Lactate dehydrogenase, LDH

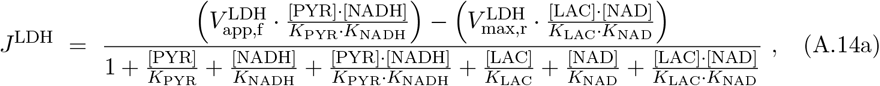

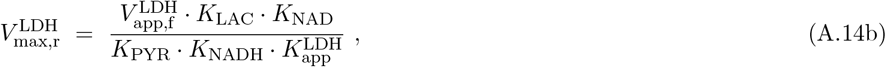

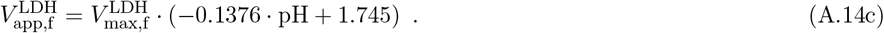

#### B.1.4 Redox buffer

##### Glycerol-3-phosphate dehydrogenase, G3PDH

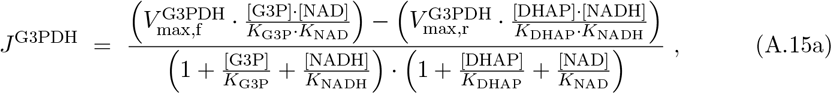

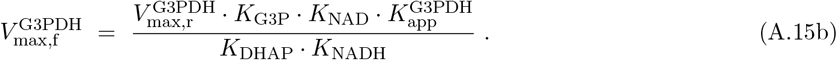

#### B.1.5 The phenomenological OxPhos model

The reference reaction of OxPhos was defined as:

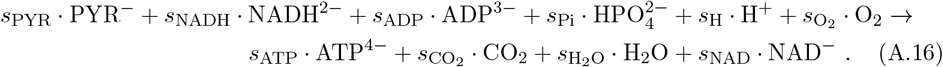

The stoichiometry is based on the following considerations. First, the respiratory quotient RQ (i.e., moles of CO_2_ produced per consumed mole of O_2_) was observed to be approximately 1.0 in resting and partially activated mouse fast twitch muscle (EDL; extensor digitorum longus) [22]. This reflects the primary breakdown of carbohydrates to generate acetyl-CoA. Thus, we only considered pyruvate as a substrate for the OxPhos model. The stoichiometry of the OxPhos reaction was calculated for the uptake of 1 pyruvate:

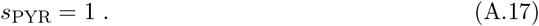

Each pyruvate consumed is coupled to the generation of reducing agents in the form of 1 NADH from the pyruvate decarboxylation, 3 NADH from the citric acid cycle (TCA), and 1 QH_2_ from the oxidation of succinate in complex II. In addition, myoplasmic NADH from glycogenolysis can be utilized in the Electron Transport Chain (ETC). Thereby, the stoichiometry factor of NADH in Eqn. (A.16) was modelled as a function of myoplasmic NADH concentration to ensure mass conservation even under extreme conditions with limited (or no) availability of myoplasmic NADH:

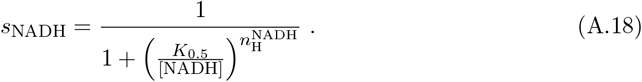

We assumed the half saturation concentration *K*_0.5_ =0.1 µM and a Hill coefficient 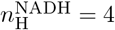. Assuming the more efficient electron transport via malate-aspartate shuttle [93], we end up with a total of (4 + *s*_NADH_) NADH and 1 QH_2_ as reducing agents. Ten protons are pumped across the inner mitochondrial membrane for each oxidised NADH at complex I compared to six protons for every two electrons donated from QH_2_ to complex II. This leads to a total of (46 + *s*_NADH_ *·* 10) protons being pumped per pyruvate. The malate-aspartate shuttle is electrogenic [94], effectively consuming *s*_NADH_ protons per pyruvate, resulting in (46 + *s*_NADH_ *·* 9) protons being pumped per pyruvate. The ATP synthase stoichiometry is 8/3 [95] plus one additional proton attributed to the transport of ATP (adenine nucleotide translocator, ANT) for a total cost of 11/3 protons per ATP. Considering the substrate-level production of 1 ATP derived from the guanosine triphosphate molecule (GTP) generated in the TCA cycle, we obtain:

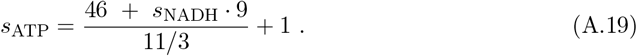

Note that we dismissed the cost of 1 proton for the transport of the additional substrate-level ATP (ANT) since the simulations were not sensitive to small changes in proton stoichiometry (result not shown). By doing so, we obtain integer values for all stoichiometry factors under normal physiological conditions, i.e. with *s*_NADH_ = 1 (see Eqn. (14)).

The oxygen consumption flux is stoichiometrically equated to half of the rate of generation of electron donors:

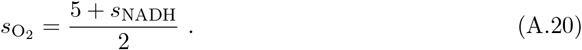

3 CO_2_ are generated per pyruvate (1 from pyruvate decarboxylation and 2 from the TCA cycle):

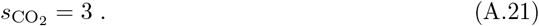

The stoichiometry factors for NAD, ADP and Pi are given by:

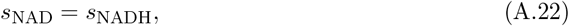

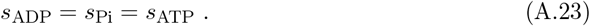

Considering the uptake of 2 H_2_O in the TCA cycle, the stoichiometry factor for H_2_O is defined as:

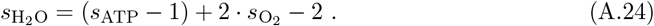

Finally, all reference reactions in the model are balanced with respect to charge, leading to:

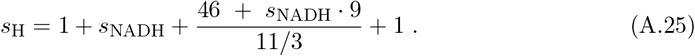

Under normal physiological conditions *s*_NADH_ equals 1 and the stoichiometry of the OxPhos model is the one given in the main text (Eqn. (14)) with a maximal P/O_2_ of 5.0 (amount of ATP produced per molecule of oxygen reduced by the ETC). With limited availability of myoplasmic NADH, the capacity for oxidative ATP production is reduced.

The kinetic equation can be found in the main text (Eqn. (15)).

#### B.1.6 Carbon dioxide/bicarbonate buffering system

##### Carbonic Anhydrase

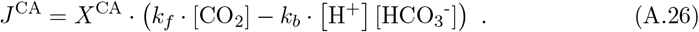

##### Uncatalyzed CO_2_ hydration (extracellular compartment)

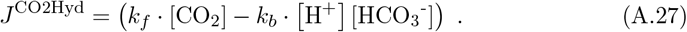

#### B.1.7 Transport mechanisms

The modelling framework includes facilitated lactate proton symport via MCT4 and passive CO_2_ diffusion between the myoplasm compartment and extracellular space. The extracellular lactate concentration was clamped to 1 mM. CO_2_ and bicarbonate are further removed from the extracellular space into a capillary compartment. In the following, [L]_my_, [L]_ex_ and [L]_cap_ indicate the concentrations of a metabolite L in the myoplasm, extracellular space and the capillary compartment, respectively. 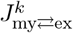 indicates transport mechanism *k* between the myoplasmic and extracellular compartment and 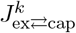 indicates transport mechanism *k* between the extracellular space and the capillary compartment. The kinetic equations of all transport mechanisms were taken from a model of glycogenolytic and oxidative energy metabolism in resting skeletal muscle fibers [81]. The diffusion parameter of 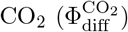 and the washout parameter of 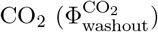 and bicarbonate transport 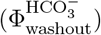 were adjusted in order to simulate actively contracting muscle fibers. The fixed CO_2_ and bicarbonate concentrations in the capillary domain 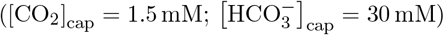 were taken from Vinnakota et al. [81]^1^. Parameter values are listed in Supplementary Table 3.

##### Monocarboxylate Transport, MCT4

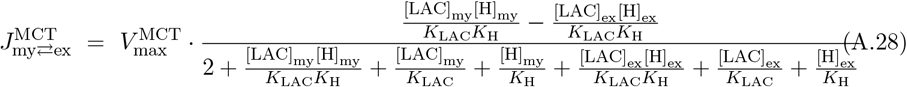

##### CO_2_ diffusion

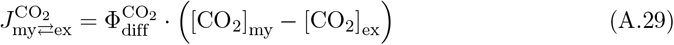

##### CO_2_ washout

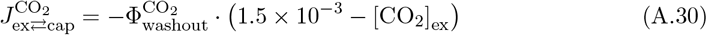

##### HCO_3_^-^ washout

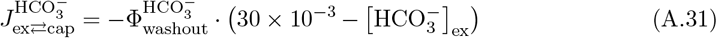

### B.2 Differential equations

The model comprises the following set of differential equations. Not included in this list are the differential equations for protons and metal ions which are obtained by the Biochemical Simulation Environment BISEN [36] as proposed by Vinnakota et al. [92]. The myoplasmic concentration of potassium, as well as the pH level in the extracellular and capillary compartments, are each clamped to the basal value reported in Supplementary Table 4 for all simulations presented in this work.

**Supplementary Table 3.** Kinetic parameters of the transport models.

| Transport model | Parameter | Parameter value |
| --- | --- | --- |
| **Myoplasmic compartment ⇌ Extracellular compartment** |  |  |
| **Monocarboxylate Transport, MCT4** | $V_{\max}^{\text{MCT}}$ | 0.088 M/min<sup>c</sup> |
| | $K_{\text{LAC}}$ | $16 \times 10^{-3}$ M<sup>b</sup> |
| | $K_{\text{H}}$ | $10^{-8.89}$ M<sup>a</sup> |
| **CO<sub>2</sub> diffusion** | $\Phi_{\text{diff}}^{\text{CO}_2}$ | 25 /min<sup>c</sup> |
| **Extracellular compartment → Capillary compartment** |  |  |
| **CO<sub>2</sub> washout** | $\Phi_{\text{washout}}^{\text{CO}_2}$ | 15 /min<sup>c</sup> |
| **HCO<sub>3</sub><sup>-</sup> washout** | $\Phi_{\text{washout}}^{\text{HCO}_3^-}$ | 1.5 /min<sup>c</sup> |

<sup>a</sup> value was taken from Vinnakota et al. [81]. Available from:
<sup>b</sup> reported $K_m$ values for lactate in skeletal muscle are between 13 mM to 40 mM for MCT4 [32, 96, 97].
<sup>c</sup> heuristically adjusted.

#### Myoplasmic compartment

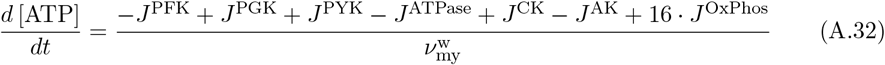

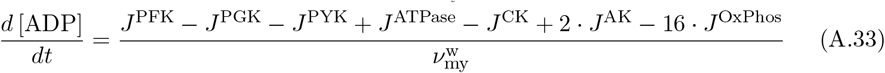

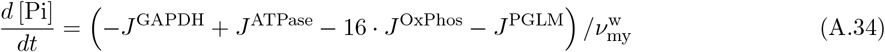

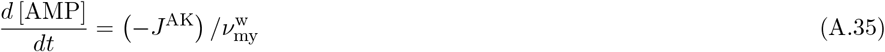

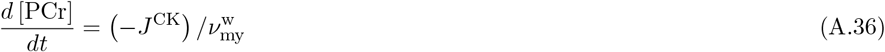

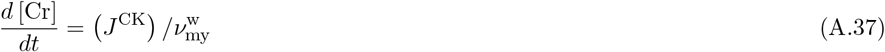

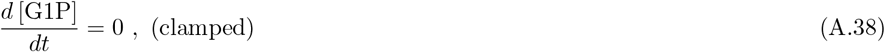

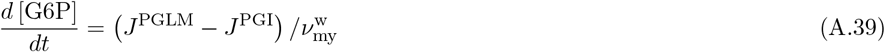

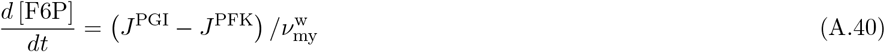

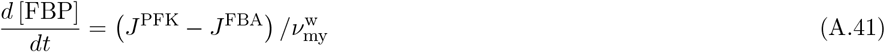

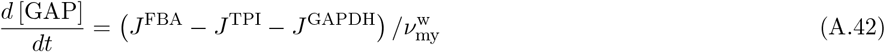

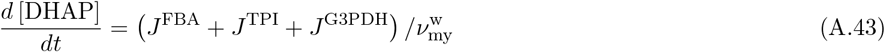

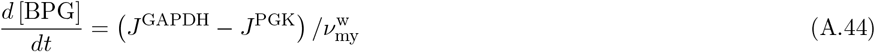

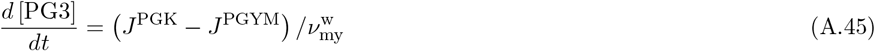

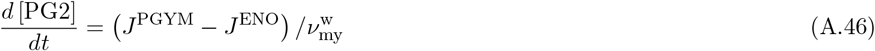

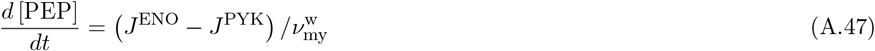

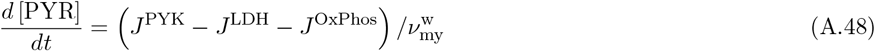

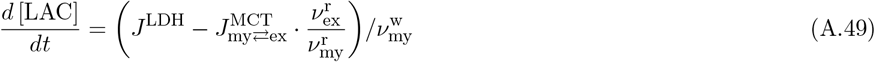

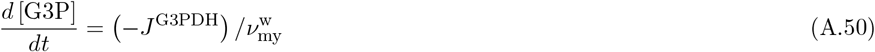

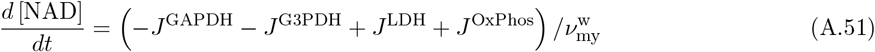

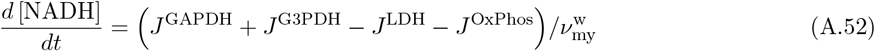

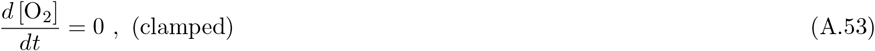

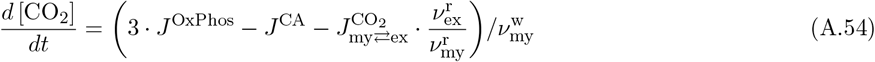

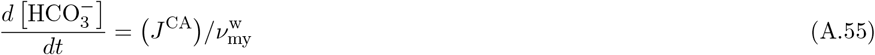

#### Extracellular compartment

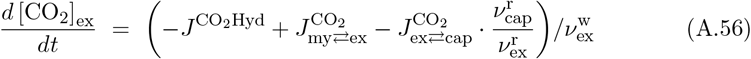

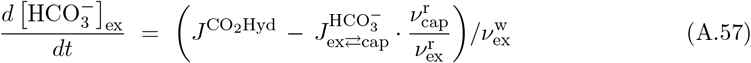

The extracellular lactate concentration is clamped per default:

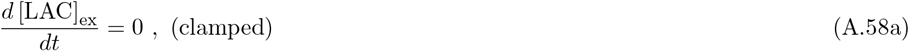

The following equation applies for the simulations under ischaemic conditions:

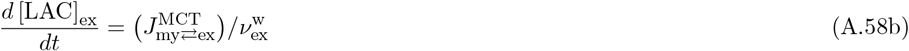

#### Capillary compartment

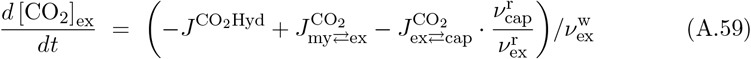

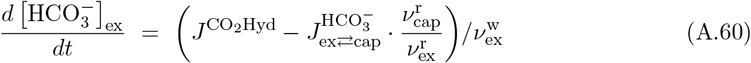

### B.3 Baseline concentrations

The resting baseline concentration of each metabolite in the model is listed in Supplementary Table 4. The table also contains reported concentrations in resting skeletal muscles from literature. However, note that the reported values are a mix of biopsy data and in vivo ^31^P-MRS data collected from different FT skeletal muscles in different species and are intended as a rough guide only.

## C MPSA

A local multi-parameter sensitivity analysis (MPSA) was conducted with the algorithm summarized in the following (for further details see [44–47]):

1. A total of 91 parameters were tested. 85 kinetic parameters, 20 of which are 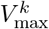 values, plus the basal pH level and 5 metabolite pools (total phosphate pool, TPP; total creatine pool, TCr; total adenine nucleotide pool, TAN; total NAD and NADH redox pool, redox; total generic buffer size, BX). The apparent equilibrium constants 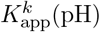 of the enzyme reactions are computed as a function of pH and were not included.
2. The perturbation range for the tested parameters was set to *±*1 %.
3. A Latin hypercube sampling method was used to generate 3000 parameter sets drawn from the given parameter range. The distributions of the parameter values in an intact physiological system are unknown, we thus assumed a uniform parameter distribution. The sampling method was implemented using the MATLAB function ‘lhsdesign’. The random number generator was initialized using the default algorithm and seed.
4. The model was run with each generated parameter vector ***p*** simulating three exercise intensities following a protocol of a 5 min resting phase with basal ATP hydrolysis rate, a 5 min exercise phase with an ATPase rate above basal level (low intensity: ATPase = 10 mM*/*min; medium intensity: ATPase = 20 mM*/*min; high intensity: ATPase = 40 mM*/*min) and a subsequent 5 min recovery phase at basal ATPase rate. An initialization phase was implemented so that the model could reach a steady-state baseline before the simulation protocol started. The pH level was clamped during the initialization phase. To evaluate the effect of a parameter perturbation on the simulation output with respect to 10 selected state variables *m* ([Pi], [PCr], pH, [ATP], [ADP], [AMP], [FBP], [G3P], [PYR] and [LAC]), a loss function was defined as the sum of squared errors between the simulation with the reference parametrization 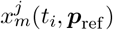 and the simulations with the randomly generated parameter sets 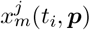:

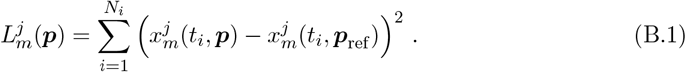

Therein, *N*_*i*_ denotes the number of time samples *t*_*i*_. The loss function value for each parameter set ***p*** with respect to the *m*^th^ selected state variable was calculated for the metabolite baseline (*j* = 1: basal ATPase rate, i.e. 0.48 mM*/*min) and the three dynamic responses, combining the exercise phase at a different ATPase rate, respectively, and the subsequent recovery phase (*j* = 2, 3, 4). For the evaluation of the dynamic response, the baseline of the reference simulation and the perturbed simulation results were shifted to zero.
5. The parameter sets ***p*** were classified to be either acceptable or unacceptable through thresholding. The mean loss function value (considering all 3000 simulations with different parameter sets) for each metabolic state variable was used as the corresponding threshold. Note that there are separate thresholds for the evaluation of the resting state sensitivity (mean loss function value over all simulation runs at rest) and for the evaluation of the dynamic response of the model at each level of muscle activation (mean loss function value over all simulation runs during the active and recovery phase combined). A parameter set ***p*** is acceptable in reference to one of the metabolic state variables (at working rate *j*) if the loss function value is below the associated threshold value and unacceptable if the value is above.
6. For each tested parameter, the cumulative distribution functions of the parameter values from the accepted and unaccepted parameter sets were evaluated using a Kolmogorov-Smirnov (K-S) statistic given as:

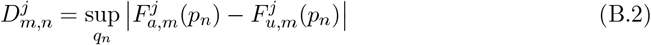

where 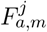 and 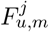 are the cumulative distribution functions of the accepted and unaccepted parameter values, respectively, of a tested parameter with respect to the *m*^th^ metabolic state variable at work rate *j*. The K-S statistic determines the largest absolute vertical difference between the two distribution functions. Consequently, if the K-S value (*D*_*m,n*_) is large, the *m*^th^ state variable is classified as sensitive to changes of the *n*^th^ tested parameter.

**Supplementary Table 4.** Basal concentrations in the model versus reported values.

| Metabolite | Model (mM)<sup>a</sup> | Reported values (mM)<sup>b</sup> | Reference |
| --- | --- | --- | --- |
| G1P | 0.07* | 0.01–0.24 | [98–102] |
| G6P | 0.2 | 0.28–0.84 | [5, 98–102] |
| F6P | 0.23 | 0.05–0.23 | [79, 98–100, 102] |
| FBP | $0.9 \times 10^{-3}$ | $40 \times 10^{-3}$–$120 \times 10^{-3}$ | [99, 100, 102] |
| DHAP | 0.055 | 0.004–0.12 | [100, 102, 103] |
| G3P | 0.59 | 0.13–0.35 | [5, 98] |
| GAP | 0.003 | 0.04 | [79, 100] |
| 13BPG | 0.003 | 0.065<sup>c</sup> | [79] |
| 3PG | $0.22 \times 10^{-3}$ | $6.6 \times 10^{-3}$–$56.3 \times 10^{-3}$ | [79, 103] |
| 2PG | $0.07 \times 10^{-3}$ | $5 \times 10^{-3c}$ | [79] |
| PEP | $0.29 \times 10^{-3}$ | $19.4 \times 10^{-3}$ | [79] |
| PYR | 0.021 | 0.022–0.17 | [5, 38, 98–100, 102, 103] |
| LAC | 0.37 | 0.36–2.23 | [5, 38, 98–101, 103] |
| ATP | 10.3 | 8.0–11.4 | [5, 37, 38, 98–102, 104, 105] |
| ADP | 0.021 | 0.009–1.79 | [5, 38, 98–100, 102, 104] |
| AMP | $0.02 \times 10^{-3}$ | $0.02 \times 10^{-3}$–0.105 | [5, 79, 98, 100, 102] |
| Pi | 3.13 | 0–4.7 | [37, 38, 102, 104, 105] |
| PCr | 34.63 | 30.1–43.9 | [5, 37, 38, 98–102, 104, 105] |
| Cr | 10.6 | 1.3–22.6 | [5, 98, 103, 104] |
| NAD/NADH | 0.85/0.002 = 425 | 294–684 | [6, 106, 107] |
| CO<sub>2, aq</sub> | 8.86 | 18.6 | [108] |
| CO<sub>2, g</sub> | 1.67 | 1.63 | [108] |
| H | $10^{-7.05}$ | $10^{-7.02}$ – $10^{-7.2}$ | [37, 38, 43, 104] |
| K | 120* | 100–168 | [109, 110] |
| Mg | 1.06 | 0.58–1 | [38, 103] |
| extracellular |  |  |  |
| LAC | 1.0* | 1.0–1.92 | [111, 112] |
| CO<sub>2, aq</sub> | 30 | - | - |
| CO<sub>2, g</sub> | 1.64 | - | - |
| H | $10^{-7.4*}$ | $10^{-7.23}$–$10^{-7.4}$ | [113] |
| K | 120 | - | - |
| Mg | 2.2 | - | - |

<sup>a</sup> M = mol/(litre cytoplasmic water). <sup>b</sup> If necessary, a factor of 3.8 [81] was used as wet-to-dry weight ratio and a factor of 0.66 [35] was used to account for the fraction of intracellular water per unit wet weight of muscle for the unit conversion into Molar. <sup>c</sup> Concentration was estimated by the order of magnitude reported for other cell types [79]. \* clamped value.

## D Supplementary material: Results

### D.1 Parameter identification: Correlation coefficients

To characterize the precision of the parameter estimation described in Section 2.2.2, we calculated the matrix of Pearson correlation coefficients. The correlation coefficients were determined using the solutions for the parameter set ***q*** from the 20 optimizations runs with different initial populations. Supplementary Fig. 2 shows the correlation matrix together with p-values.

### D.2 Experiment 1: Redox contributions, pyruvate and lactate dynamics

The proposed metabolic model contains four reactions contributing to the cytoplasmic redox state. Their total effect on the redox state is summarized in Fig. 5. Supplementary Fig. 3 shows for three contraction intensities (low, medium, and high) the flux rates through each individual reaction contributing to the myoplasmic redox state, i.e., the reactions catalysed by GAPDH, G3PDH, and LDH as well as OxPhos. The simulations showcase a tightly balanced system of NAD^+^ reduction and NADH oxidation, enabling rapid changes in metabolic flux rates (in the range of mM*/*min) yielding considerably lower total NADH flux rates in the micro Molar per minute range (see Supplementary Fig. 3D-F). Notably, the redox potential reaches a steady state in less than two minutes after the onset of muscle activation, although several reactions still show transient behaviour for example the G3PDH-reaction.

**Supplementary Fig. 2.**
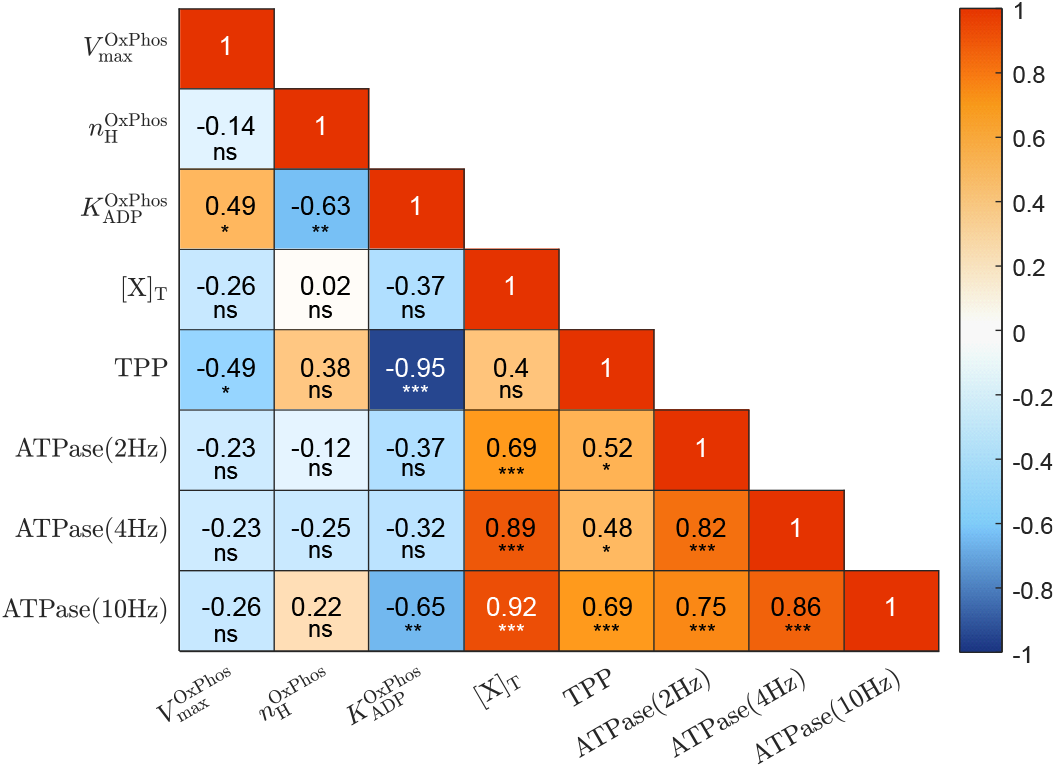
Correlation matrix of optimized parameter values. The significance of the correlation is indicated by symbols (ns : *p*-value ≧ 0.05, ^*\**^ : *p*-value *<* 0.05, ^*\*\**^ : *p*-value *<* 0.01, ^*\*\*\**^ : *p*-value *<* 0.001).

Further, the intracellular lactate and pyruvate concentration dynamics are shown in Supplementary Fig. 4 for the same simulation protocol. Lactate accumulation as well as a comparatively small accumulation of pyruvate are observed for all three simulated contractions intensities, including low intensity exercise, with ATPase rates below the maximal capacity for oxidative ATP production (Supplementary Fig. 4A). An accumulation of both lactate and pyruvate facilitate an increased pyruvate consumption through OxPhos. Thus, in a biochemical network including LDH, lactate accumulation during low intensity exercise is a necessary prerequisite to steer pyruvate uptake in the direction of OxPhos through increased levels of PYR.

### D.3 Experiment 1: Transformed Gibbs energy of the redox half reaction

The biochemical reaction of the redox half reaction is given by:

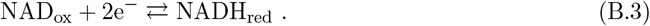

The associated reference reaction is defined as:

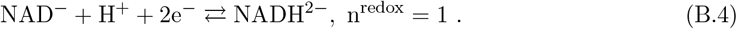

The redox half-reaction is not a stand-alone reaction in the model. Thus, the apparent equilibrium constant necessary for the calculation of the transformed Gibbs energy of the biochemical reaction (see Eqn. (11)) is not explicitly computed as part of the model. Instead, the value was calculated separately using the same method as implemented in BISEN. We used the standard Gibbs energies of formation listed in the BISEN database (see Table 5).

**Supplementary Fig. 3.**
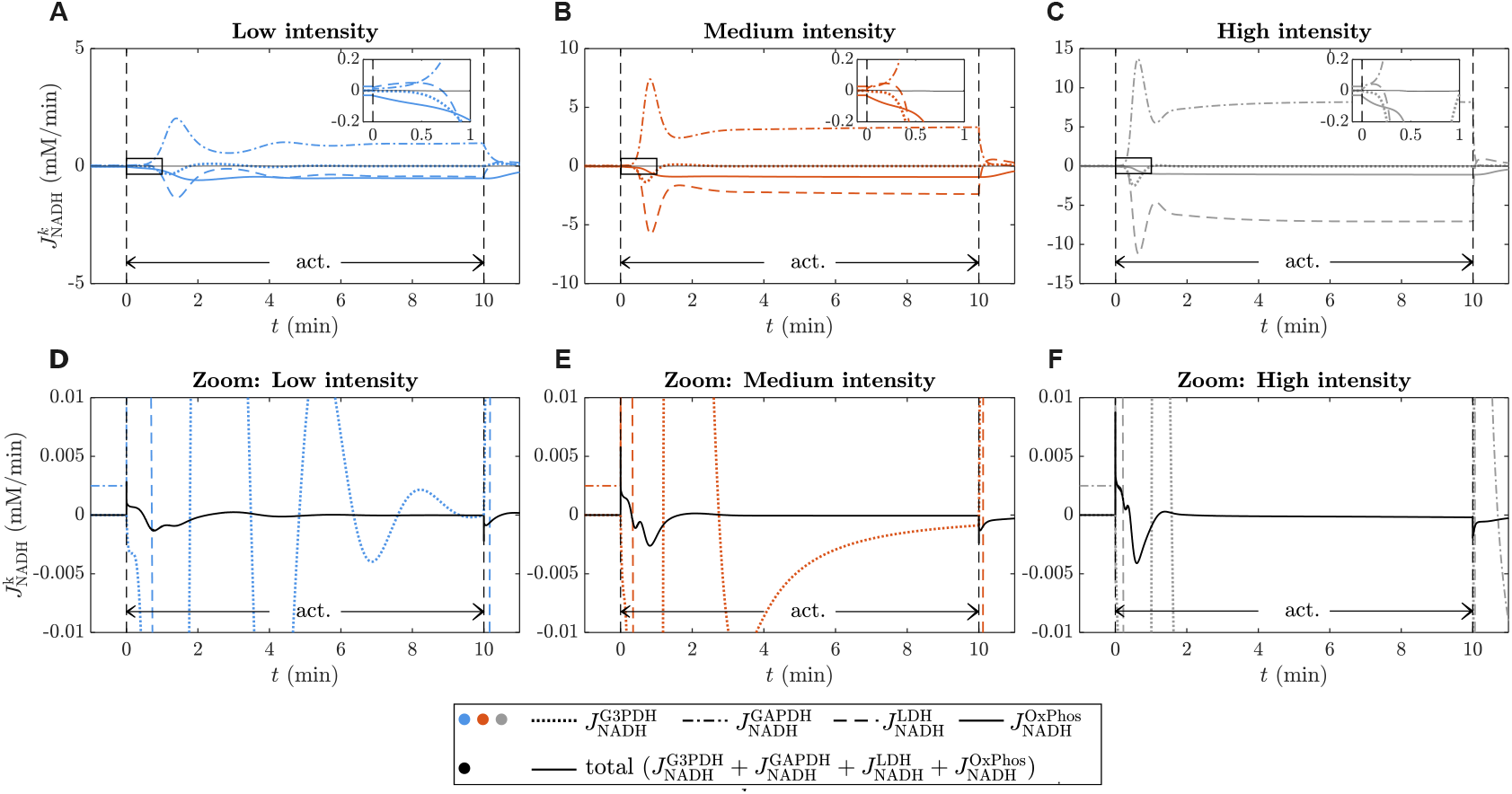
NADH flux rate 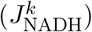 through each associated (enzyme-catalysed) reaction for three different contraction intensities (Blue: ATPase rate 10 mM*/*min, Orange: ATPase rate 20 mM*/*min, Gray: 30 mM*/*min). A positive flux indicates NADH generation, a negative flux indicates NADH oxidation.

**Supplementary Fig. 4.**
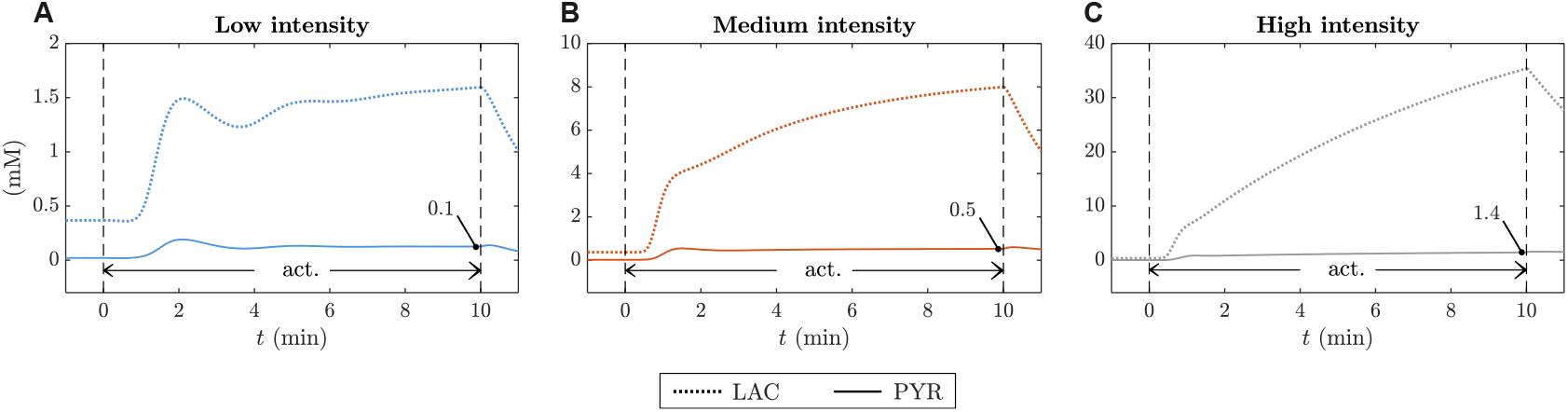
Pyruvate and lactate time course during 10 min muscle exercise for three different contraction intensities, i.e., ATPase rate 10 mM*/*min (A), ATPase rate 20 mM*/*min (B), ATPase rate 30 mM*/*min (C)

**Supplementary Table 5.** Properties of the reference species of the redox half reaction.

| Reference species | $\Delta_f G^0$ (kJ/mol)<sup>a</sup> | $v_i$ | $z_i$ |
| --- | --- | --- | --- |
| H<sup>+</sup> | 0 | -1 | 1 |
| NAD<sup>-</sup> | 0 | -1 | -1 |
| NADH<sup>2-</sup> | 22.65 | 1 | -2 |

<sup>a</sup>values were taken from the BISEN data base [36] (see also [11]). The values are determined at 25°C, 1 Bar and I = 0.

The standard Gibbs energy of the reference reaction is the difference in the energies of formation of the reactants and products:

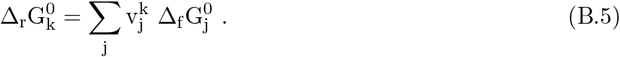

The Gibbs energies of formation of the reference species in Supplementary Table 5 were reported at zero ionic strength. The standard Gibbs energy of the reference reaction was empirically adjusted to the ionic strength in the model using the following equations:

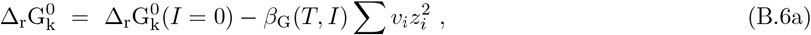

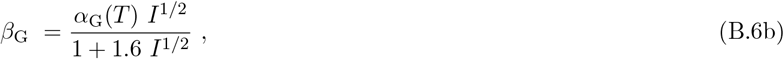

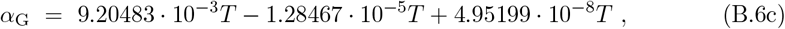

with *I* = 0.1 M and *T* = 298.15 K representing the ionic strength and temperature used in the presented simulations. The stoichiometry factors, *v*_*i*_, and charges, *z*_*i*_, of the reference species are listed in Supplementary Table 5. We note that the BISEN data base does not include any enthalpies of formation necessary for the adjustment of Δ_r_G^0^ to the temperature in the model even though literature values are available [11]. We decided to forgo the empirical temperature correction to stay consistent with the rest of the model.

The equilibrium constant of the reference reaction was calculated (Eqn. (10)) in order to compute the apparent equilibrium constant for the associated biochemical reaction using the simulated proton dynamics (Eqn. (8)). The dissociation constant for the binding polynomials (Eqn. (9)) are not specified in the BISEN data base for the metabolites involved and, therefore, were assumed to be infinite (no binding).

Finally, *K*_app_ was used to determine the the dynamics of the transformed Gibbs energy of the redox half reaction (Eqn. (12) and (11)).

### D.4 Experiment 2: Redox contributions, pyruvate and lactate dynamics

The conducted in silico experiment 2 compares the behaviour of the control model and the LDH-KO model given a 20-minute-long exercise protocol with ATPase rates up to the maximal rate of oxidative ATP production 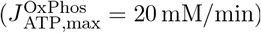. Supplementary Fig. 5A-B exemplarily shows the NADH flux rates through the reactions catalysed by GAPDH, G3PDH and LDH as well as OxPhos during low intensity and medium intensity exercise. Lactate and pyruvate dynamics for the same simulation protocol are shown in Supplementary Fig. 5C-D. Supplementary Fig. 6 (A and D) compares the NADH time course of the control model and the LDH-KO model for a selection of simulated ATPase rates ranging from 0.5 mM*/*min to 20 mM*/*min. The simulation results suggest that LDH activity dampens oscillations in NADH content during jumps in ATP demand at ATPase rates. The transient limited availability of myoplasmic NADH in the LDH KO system at some of the simulated exercise intensities results in a decrease in myoplasmic NADH utilization by the aerobic pathway (Supplementary Fig. 6C,F) and thus, a reduced capacity for oxidative ATP production (Supplementary Fig. 6B,E). The observed change in the ATP (*s*_ATP_) and NADH (*s*_NADH_) stoichiometry factor of the OxPhos model is very small, however, necessary to avoid complete NADH depletion and to ensure mass conservation (see Appendix B.1.5).

**Supplementary Fig. 5.**
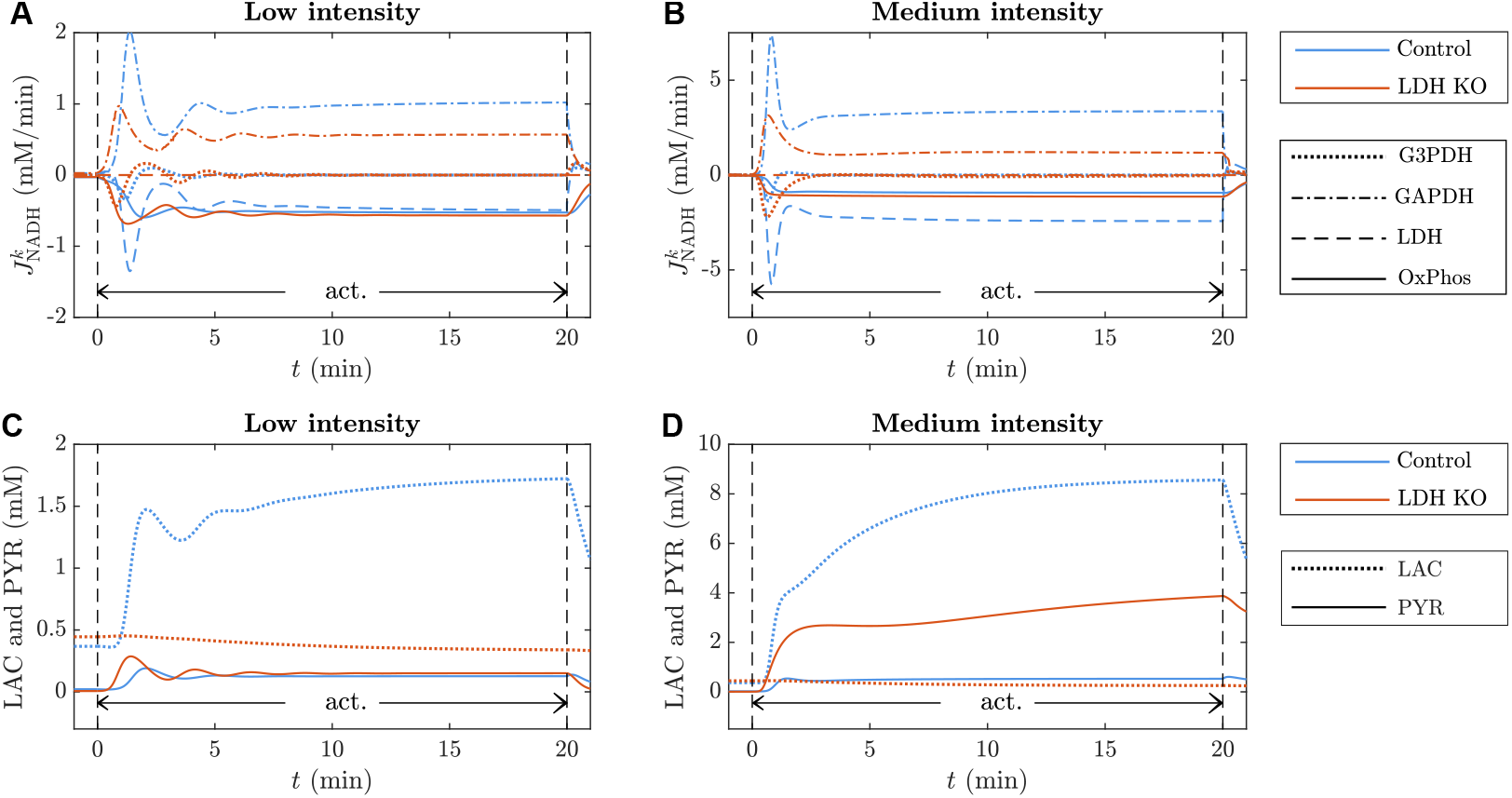
Comparison of the control system (blue) and LDH KO system (orange) during 20 min muscle exercise at two different contraction intensities (low intensity: ATPase rate 10 mM*/*min, medium intensity: ATPase rate 20 mM*/*min). (A-B) NADH flux 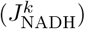 through each associated (enzyme-catalysed) reaction. A positive flux indicates NADH generation, a negative flux indicates NADH oxidation. (C-D) Pyruvate and lactate time course.

### D.5 Underdamped dynamic behaviour

The model predicts a dynamic behaviour similar to a second-order underdamped system in transitions between steady states, prominently at low ATP turnover rates. This behaviour is manifested, for example, in the time course of myoplasmic PCr and NADH concentrations (Supplementary Fig. 7A,D) reflecting the oscillating flux rates of the biochemical reactions in the metabolic network. Supplementary Fig. 7G exemplarily shows the metabolic fluxes associated with NADH concentration in the muscle fiber. To investigate the underlying mechanisms of this behaviour, we simulated the dynamic response to 10 min of low intensity exercise (ATPase rate: 10 mM*/*min) of the control model, a model configuration with increased 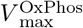 and a model configuration with increased 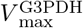. Increasing 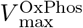 to 1.5-times the control value enhances the underdamped behaviour of the system with oscillations lasting throughout the entire 10 min of muscle activation (Supplementary Fig 7B,E,H). In contrast, increasing 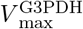 to double its control value dampens the oscillations in metabolic fluxes and, thus, the myoplasmic PCr and NADH concentrations slightly compared to the control system (Supplementary Fig 7C,F,I).

**Supplementary Fig. 6.**
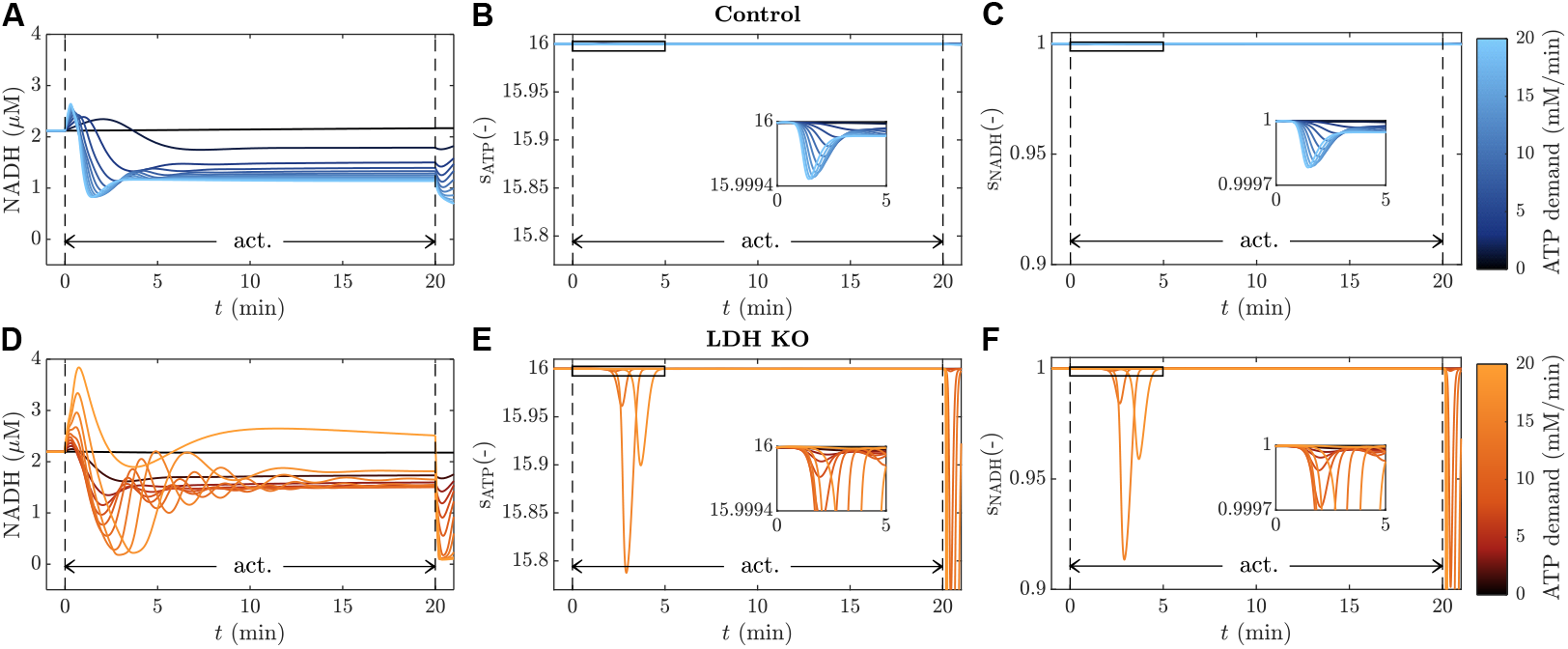
Comparison of the control system (A-C) and LDH-KO system (D-F) during 20 min muscle exercise at a selection of 12 ATPase rates ranging from 0.5 mM*/*min to 20 mM*/*min. The ATPase rate is indicated by the colour gradient. (A and D) NADH dynamics. (B and E) dynamic change in OxPhos stoichiometry factor for ATP and (C and F) dynamic change in OxPhos stoichiometry factor for NADH.

**Supplementary Fig. 7.**
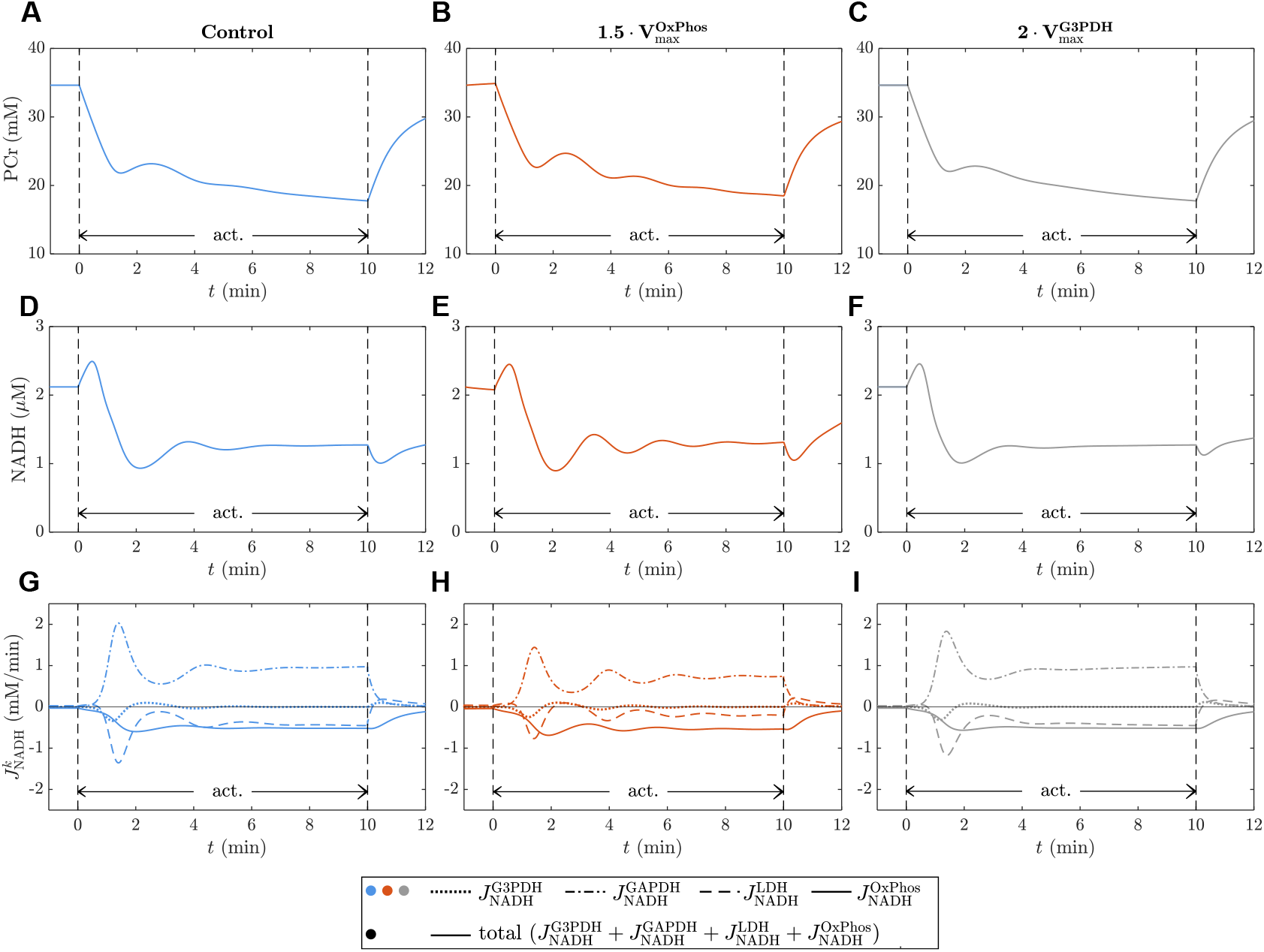
Comparison of the dynamic response of the control model (left), a model configuration with increased 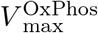 (middle) and a model configuration with increased 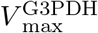 (right) to 10 min of muscle activation at an ATPase rate of 10 mM*/*min. (A-C) PCr concentration. (D-F) NADH concentration. (G-I) NADH flux 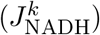 through each associated (enzyme-catalysed) reaction. A positive flux indicates NADH generation, a negative flux indicates NADH oxidation.

## Acknowledgments

This research was funded by the German Research Foundation (Deutsche Forschungsgemeinschaft, DFG) as part of the cluster of excellence “Data-integrated simulation science (SimTech)” (EXC 2075 390740016) and the priority program SPP 2311 (465195108 and 548605919), the European Research Council (ERC) through ERC-AdG ‘qMOTION’ (101055186), the National Institutes of Health (NIH) through grants HL173346 and HL154624 and by Stichting Spieren voor Spieren of the Netherlands.

## Data availability statement

The source code and data used to generate the results and analyses in this study are openly available at the following URL: https://github.com/j-disch/EnergyMetabolism_FOG.git.

## Footnotes

1 Available from: https://models.physiomeproject.org/exposure/aa821a8dda7e55888bd486f8cbb68791/vinnakota_rusk_palmer_shankland_kushmeric_2010_edl.cellml/view; Accessed: 2025-01-15; CellML author: Geoffrey Nunns.

